# Dynamics of fluctuating populations in multi-state switching environments

**DOI:** 10.64898/2026.08.11.744206

**Authors:** Mauro Mobilia

## Abstract

Microbial populations generally evolve in fluctuating environments under time-varying conditions. These are often described by binary switching models, sometimes seen as coarse-grained feast–famine cycles, in which resource availability switches abruptly between abundant and scarce conditions. However, experimental studies suggest that feast–famine environments actually exhibit more complex temporal dynamics. Here, we study how two strains, one growing slightly slower than the other, compete for the same resources in fluctuating environments comprising a finite number of intermediate states, each having its own carrying capacity. Environmental switching between these states and their carrying capacities represents gradual changes in nutrient availability. This class of multi-state stochastic switching models can be interpreted as a coarse-grained description of feast–famine cycles and allows us to investigate strain competition under the gradual recovery and depletion of resources. By computational and analytical means, we characterise the population dynamics in these multi-state fluctuating environments. In particular, we study how the switching rates and distribution of carrying capacities affect the population-size statistics, fixation probability, and mean fixation time. By comparing these results with their counterparts in binary environments, we clarify how the frequency and amplitude of environmental fluctuations influence population dynamics in coarse-grained feast–famine cycles.

## I. INTRODUCTION

Microbial populations live in volatile and fluctuating environments in which available resources, as well as temperature, pH, toxin concentration, can fluctuate substantially over time [1–3]. These abiotic variations, referred to as environmental fluctuations (EF), can have an important influence on the evolution of biological populations [4–7]. In small populations, demographic fluctuations (DF) are another important form of randomness and can result in fixation when one type takes over the population - or extinction [8–10]. The variations of population size and composition are often interdependent [11– 17], and this can lead to the coupling of EF and DF [18–28]. The interplay of environmental variability and DF is particularly relevant when it gives rise to population bottlenecks, where new small microbial colonies are prone to fluctuations [29–31]. It has recently been shown that environmental variability, bottlenecks and fluctuations are particularly relevant for the dynamics of antimicrobial resistance [3, 24, 26, 28, 32–35]. The influence of different kinds of variability, such as heterogeneous rates and interactions, on the eco-evolutionary dynamics of competing species has also received much attention [36–38]. Population dynamics in fluctuating environments have traditionally been studied both theoretically and experimentally using two-state, or binary, environmental switching [7, 39–46], where individuals compete for resources under two alternating environmental conditions. For instance, in Refs. [18–22, 24–28], the time-varying environment is represented by a binary switching carrying capacity, whose fluctuations drive the population size. This in turn modulates demographic noise, leading to the coupling of EF and DF, which has a significant impact on the population size distribution and its fate, and is characteristic of cycles of abundant and scarce resources.

Feast–famine cycles refer to alternating periods of nutrient abundance (“feast”) and resource deprivation (“famine”) experienced by many microbial populations [47–50]. The temporal statistics of these nutrient fluctuations strongly influence long-term microbial adaptation by mediating the trade-off between growth during resource-rich periods and survival during starvation, thereby shaping the evolution of different ecological strategies, including generalist and specialist nutrient-utilization strategies [47, 48, 50]. Competition between strains in an environment with a binary time-switching carrying capacity can be viewed as the simplest coarse-grained representation of a feast–famine cycle, where the two possible values of the carrying capacity correspond to the environmental states of feast and famine [18– 21]. While this binary switching model has the advantage of being mathematically tractable, recent studies have highlighted the importance of multiple intermediate environmental conditions in population dynamics subject to feast–famine cycles. For instance, the coupled nutrient and population dynamics considered in Ref. [49] naturally generate intermediate conditions between feast and famine, which are shown to play an important dynamical role. Related work has further investigated the evolutionary consequences of fluctuating nutrient availability for competing generalist and specialist nutrient-utilization strategies under stochastic resource supply [50]. More generally, experiments have demonstrated that the temporal statistics of EF, in particular their frequency and amplitude, strongly influence microbial population dynamics under feast–famine cycles [48]. Furthermore, the experimental comparison of eco-evolutionary dynamics in binary and ternary environments revealed important dynamical differences between abrupt environmental switching and gradual environmental deterioration through intermediate conditions [51]. These studies therefore suggest that feast– famine environments possess a richer temporal and statistical structure than can, in general, be captured by binary switching models. This motivates us to investigate stochastic multi-state environmental models with intermediate environmental states characterized by distinct carrying capacities.

Specifically, here we study the dynamics of two strains, one slightly slower than the other, competing for the same resources in multi-state fluctuating environments. The time-varying environment consists of multiple intermediate states, ordered from the harshest (“famine”) to the mildest (“feast”) conditions. Each environmental state is associated with its own carrying capacity, whose value increases progressively from harsh to mild states; see Fig. 1 (a). We therefore consider a class of multi-state stochastic switching models that provide a coarse-grained representation of feast–famine cycles with intermediate environmental states. These stochastic individual-based models allow us to describe, at population level, the strain competition when the recovery and depletion of resources is gradual [48–51]. This is an important feature of feast–famine cycle dynamics that is not captured by binary models. To characterise the multi-state switching dynamics, we ask: *How do the environmental switching rates and the distribution of carrying capacities influence the population size statistics, as well as the probability and mean time of fixation? How do these quantities differ from their binary-state counterparts?*

**FIG. 1.**
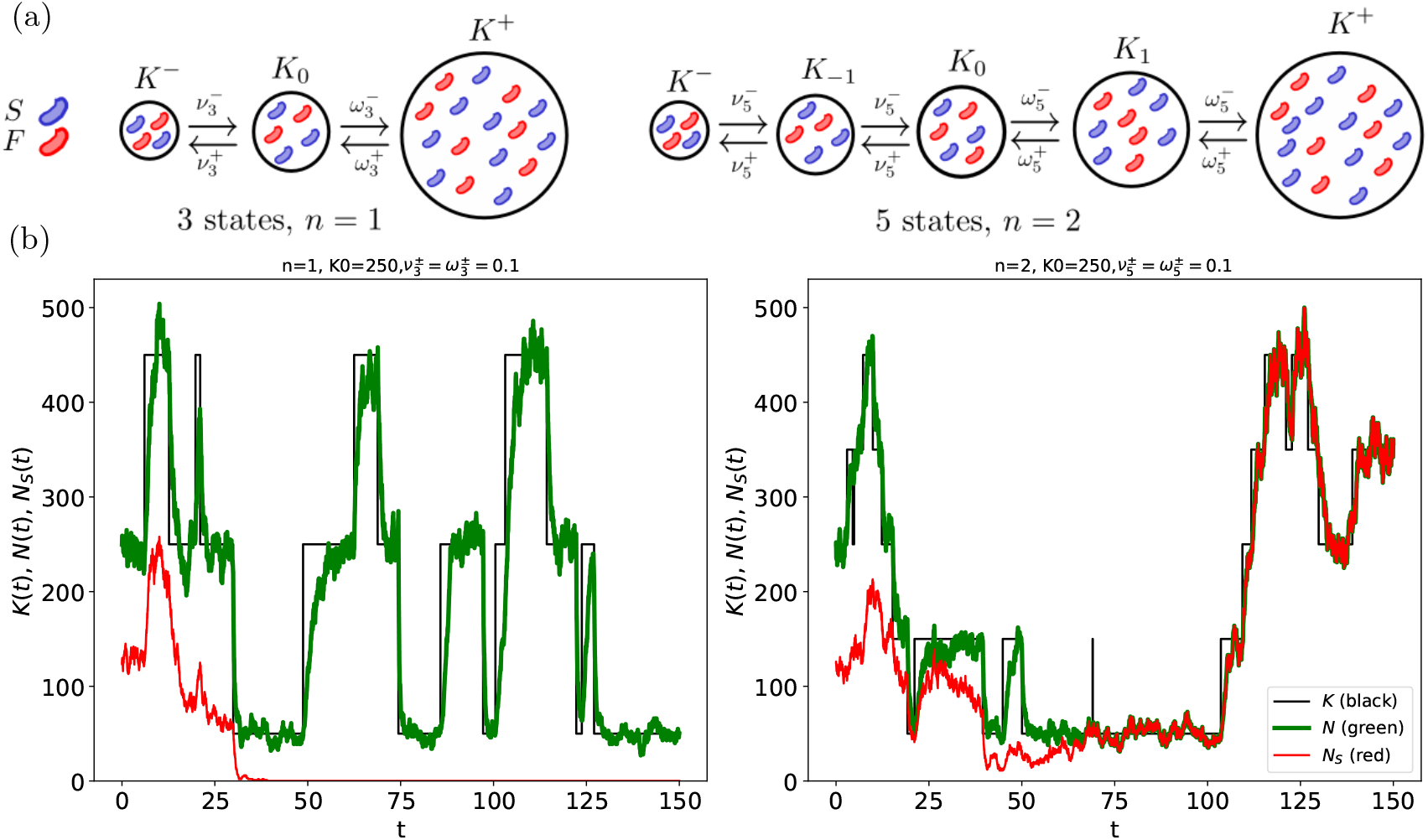
(a) Illustration of the the 3-state (*n* = 1, left) and 5-state (*n* = 2, right) switching eco-evolutionary dynamics.(b) Typical sample paths of *K*(*t*) (black), *N* (*t*) (green), and *N*_*S*_ (*t*) (red) for the 3-state (left) and 5-state (right) switching models with (*K*^−^, *K*^+^) = (50, 450). The number of *F* individuals is *N*_*F*_ (*t*) = *N* (*t*) − *N*_*S*_ (*t*) (not shown). (b, left) *n* = 1, *K*_0_ = 250, *K*(*t*) ∈ {50, 250, 450}, 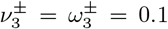 (intermediate switching regime; see text); (b, right) *n* = 2, *K*_0_ = 250, *K*(*t*) ∈ {50, 150, 250, 350, 450}, 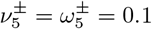 (intermediate switching). In (b, left), simultaneous *S* extinction and *F* fixation (not shown) occurs at *t* ≈ 35, whereas in (b, right) there is *S* fixation (with extinction of *F*, not shown) at *t* ≈ 70. After the fixation of a strain and extinction of the other, the population consists of only the fixed type (*F/S* on the left/right panel) and, driven by the switches of *K*(*t*), *N* (*t*) keeps fluctuating. Other parameters in (b) are (*s, x*_0_) = (0.02, 0.5).

We address these questions by using simulation and analytical means to compute these quantities and compare them in multi-state and binary switching models. Since it has been experimentally shown feast–famine cycle dynamics depend strongly on the frequency and amplitude of EF, we particularly study the influence of the switching rates (EF frequency) and distribution of carrying capacities – encoding the EF amplitude – on the population size distribution and on its fixation properties. This sheds further light on how the frequency and amplitude of environmental fluctuations shape the complex dynamics of coarse-grained feast–famine cycles.

In the next section, we introduce the class of multistate feast–famine models that we study and outline our methods. In Sec. III, we present our results, first for the population size distribution and average population size (Sec. III A) and then discuss in detail the influence of the environment on the fixation probability of the slow strain (Sec. III B). Sec. IV is dedicated to a discussion of our findings and to our conclusions. Additional technical details, as well as supplementary results, are given in a series of appendices.

## II. MODELS & METHODS

We consider a well-mixed population of time-fluctuating size N(*t*) = *N*_*S*_(*t*) + *N*_*F*_ (*t*) that at time *t* consists of *N*_*S*_ individuals of the slow growing strain *S*, and *N*_*F*_ of a faster growing type *F*. Each *F* individual has a baseline fitness *f*_*F*_ = 1, while all slow growers have fitness *f*_*S*_ = 1− *s*. We assume that *S* and *F* compete for the same resources, with population growth limited by the carrying capacity denoted by *K*. The fraction of slow growers in the population is *x* = *N*_*S*_*/N*, and the average population fitness is thus 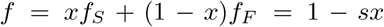. Therefore, the per capita growth rates of *F* and *S* are respectively 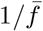 and 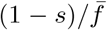, with the growth of both strains limited by a logistic death rate *N/K* [11, 13, 18, 19, 21, 23, 27, 52]. The population dynamics evolve in continuous time. For analytical tractability, we often assume 0 < s ≪1, giving a small selective advantage to *F* over *S* and yielding a time scale separation [18–21, 23, 27]; see below. However, many results of Sec. III B remain valid for 0 < s < 1.

In close relation to the Moran process [8, 53–56] (see Sec. II C), a reference model in mathematical biology, the competition (selection) dynamics is represented by a multivariate birth-death process [8, 52] defined by the birth (or division) and death of an individual of type *α* ∈ {*S, F*} according to [18–23]

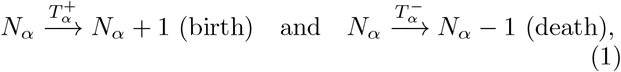

occurring at the transition rates

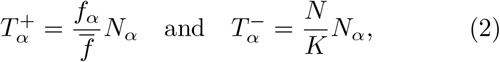

where the carrying capacity *K* is constant in a static environment and is a randomly time-varying quantity in fluctuating environments; see below. (For notational simplicity we have dropped the time dependence from 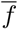 and transition rates). In this formulation, selection operates on birth events [18–28, 57–59]. This can be generalized to include selection on deaths; see e.g. Refs. [12, 13, 60].

### A. Multi-state environmental dynamics

The carrying capacity provides a coarse-grained description of environmental limitations on population growth [11], and is closely related to chemostat setups that are commonly used in laboratory-controlled experiments [3, 7, 22, 42, 61, 62]. A simple and biologically relevant way to represent environmental variability and to capture the interdependence of demographic and EF therefore consists of letting the carrying capacity be a randomly time-varying quantity [18–28]. Environmental variability is thus often described by binary-state models [7, 39–41], with the carrying capacity typically taking two possible values describing sudden and radical changes of nutrient resources [18–22, 24–28]; see Appendix A 1. Here, we introduce a class of multi-state switching models where, in addition to extreme abundance (feast) and scarcity (famine), there are 2*n* − 1 intermediate states (*n* ≥1), each with its own carrying capacity; see Fig. 1(a). The switching between the (2*n* + 1) environmental states and their carrying capacities can hence be viewed as describing a coarse-grained multi-state feast–famine cycle, with the switching rate and distribution of the carrying capacities encoding the frequency and amplitude of nutrient fluctuations [47–50]. Although it does not explicitly model nutrient dynamics, this formulation provides an individual-based description of the gradual degradation and recovery of environmental conditions that is characteristic of feast–famine dynamics [48–50].

In this context, environmental variability is encoded by the continuous-time multi-state coloured noise *ξ*(*t*) = *I* ∈ {− *n*, …, 0, …, *n*}; see Appendix A 2 that is a (2*n* + 1)state generalisation of the classical binary telegraph (dichotomous) noise [18–20, 22, 24–28, 41, 43, 63–65]. Each of the possible values *ξ*(*t*) = *i* corresponds to a specific environmental state with its own carrying capacity *K*(*ξ*(*t*)) = *K*_*i*_. The states *i* = *n*, …, *n* are ordered from the harshest (“famine”) to the mildest (“feast”). The famine and feast states are labelled by −*n* and *n*, respectively, and have carrying capacities *K*_−*n*_ ≡ *K*^−^ and *K*_*n*_ ≡ *K*^+^. (The notation *K*^*±*^ is introduced to facilitate the comparison with the binary case.) We respectively refer to *i* ∈{1− *n*, …, 1} and *i* ∈ {1, …, *n* – 1} as the intermediate harsh and mild states, while *i* = 0 is the “median state” (or “centre state”; see below) and its carrying capacity is *K*_0_. The carrying capacities are ordered and increase from harsh to mild states according to

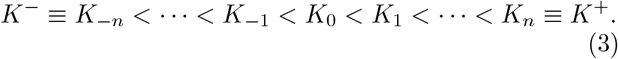

when *K*_0_ = (*K*^+^ +*K*^−^)*/*2, *i* = 0 is called the centre state. The transitions between the environmental states *i* and *i* + 1 occur with rates 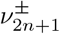 for harsh states (*i* = −*n*, …, −1), and with rates 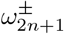 for states *i* =1, …, *n* − 1, according to

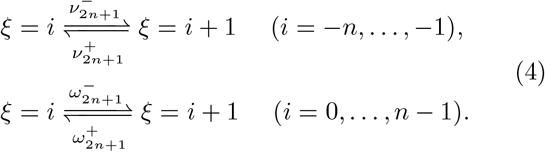

The carrying capacity 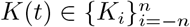 is thus a random variable whose value switches at rates 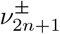 and 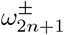 according to

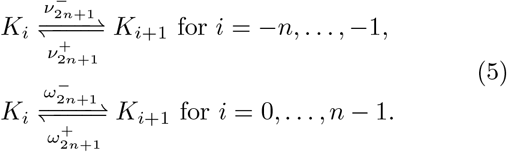

When 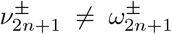, transitions among harsh states and among mild states occur at different rates. For the sake of simplicity, we introduce the parameter *ϵ* > −1 and henceforth assume 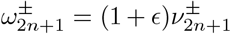. Therefore, the transitions between states *i* > 0 occur at a higher rate than those between states *i* < 0 when *ϵ* > 0, whereas it is the opposite when −1 < *ϵ* < 0. The only physical constraint on the values of *K*_*i*_ is given by the ordering relation (3). While various choices are possible, for simplicity, we consider 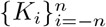 to be uniformly (linearly) distributed in [*K*^−^, *K*_0_] and [*K*_0_, *K*^+^], with

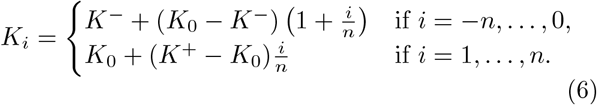

When *n* = 1 we thus have *K* ∈ {*K*^−^, *K*_0_, *K*^+^}, while in the five-state case (*n* = 2), *K* ∈ {*K*^−^, (*K*_0_ + *K*^−^)*/*2, *K*_0_, (*K*_0_ + *K*^+^)*/*2, *K*^+^}; see Appendix A 2 and Fig. 1(a). Hence, the spacing between consecutive carrying capacities in 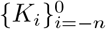 is smaller than that in 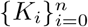 when *K*_0_ < (*K*^+^ + *K*^−^)*/*2, whereas the converse holds when *K*_0_ > (*K*^+^ + *K*^−^)*/*2. When *K*_0_ = (*K*^+^ + *K*^−^)*/*2, 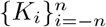 are uniformly spaced between *K*^−^ and *K*^+^, and *i* = 0 corresponds to the “centre state”.

To meaningfully compare the environmental variability in multi-state and binary models, we require that the effective binary switching rates 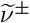 of the transitions 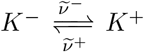 [18–21] between the e xtreme carrying capacities are such that the mean times for these transitions be the same as under multi-state switching. We notice that in (2*n* + 1)-state switching models, the transitions *K*^∓^ → *K*^±^ consists of *n* moves, each of mean waiting time 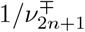, and *n* other transitions of expected waiting time 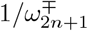 each. Matching the mean times of the transitions *K*^∓^ → *K*^±^ under binary and multi-state switching thus yields (see Appendices A 1 and A 2)

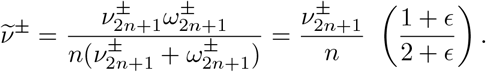

The effective rates 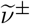 allow us to compare the environmental dynamics across binary and (2*n* + 1)-state switching models. It is convenient to write 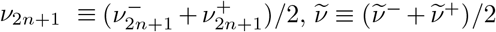 and characterise the bias between forward (*K*_*i*_ → *K*_*i*+1_) and backward switching (*K*_*i*+1_ → *K*_*i*_) by 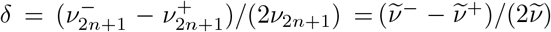 [21, 23, 24, 26, 28], with |*δ*| < 1. We thus have

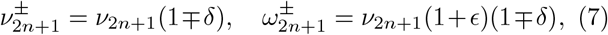

and the effective binary rates 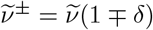, where

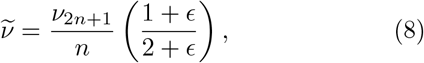

with transitions from states *i* → *i* + 1 occurring at a higher rate than those from *i* + 1 → *i* when *δ* > 0 (*i* ≠ 0). This means that *δ* > 0 corresponds to spending on average more time in states *i* + 1 than *i* (*i ≠* 0).

In this work, environmental noise and the carrying capacity are always at stationarity, with their distribution denoted by ***π*** = (*π*_*i*_) and characterised by the parameters *n* and *ρ* ≡ (1 + *δ*)*/*(1 − *δ*). In Appendix A 2, we show that

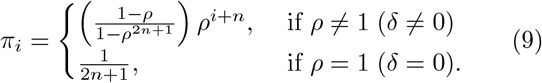

This distribution is a function of *n* and *δ*, but is independent of *ϵ* (see below). Since *π*_*i*_*/π*_−*i*_ = *ρ*^2*i*^, mild states (*i* > 0) are more likely than harsh states (*i* > 0) when *ρ* > 1 (*δ* > 0) and conversely when *ρ* < 1 (*δ* < 0). The distribution *π*_*i*_ = 1*/*(2*n*+ 1) is uniform when *δ* = 0 (symmetric forward/backward switching) [18–20, 22, 23, 27]. Here, *ξ* and *K* are always at stationarity, and their initial value in each simulation run is drawn from ***π*** according to Eq. (9); see Appendix C.

In this study, environmental dynamics is thus represented by a class of multi-state switching models, inspired by chemostat systems [3, 7, 22, 42, 61, 62], characterised by the parameters {*n, K*^±^, *K*_0_, *ν*_2*n*+1_, *δ, ϵ*}. These multistate individual-based models, defined by Eqs. (1)-(6), satisfy the master equation (A7), and are simulated using the Gillespie algorithm [66]; see Appendix C.

### B. Mean-field dynamics in a static environment

It is useful to consider the dynamics in a static environment, where the carrying capacity is constant 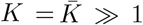. In this setting, after a transient, the population size is well approximated by the constant carrying capacity, 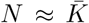 [18–22, 24–28]. When *N* and 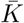 are very large, demographic fluctuations are negligible and a mean-field description of the dynamics is in order. Here, it is defined by the following rate equations for *N* and *x* [18–21]:

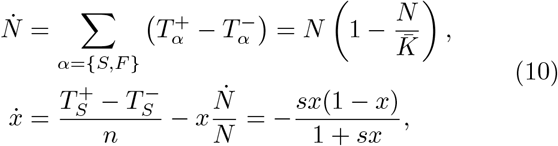

where the dot indicates the time derivative and we have used the expressions of 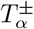 given by Eqs. (2), with *f*_*S*_ = 1 − *s* and *f*_*F*_ = 1 and 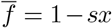 [18–21]. The logistic rate equation for *N* leads to the population size reaching the carrying capacity, 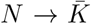, on a timescale *t* ~ 1. Since 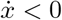 for 0 < *x* < 1, the fraction of *S* cells *x* decreases on a slower timescale *t* ~ *O* (1*/s*), and eventually vanishes (*x* → 0). We notice that, when *s* ≪ 1, there is a timescale separation, with *N* and *x* acting respectively as the fast and slow variables. Further, Eqs. (10) show that the dynamics of *N* and *x* decouple at mean-field level within a static environment. As discussed below, in this class of switching models, eco-evolutionary coupling arises from fluctuations at the individual-based level.

### C. Finite population in a static environment: Moran model

It is also relevant to consider a finite population of constant size 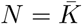 in a static environment, and study its fixation properties. This is achieved in terms of the Moran model [8, 53], which is a reference evolutionary process [55, 56, 67].

In a static environment where the carrying capacity is finite and constant, with 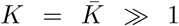, the population is unlikely to go extinct in an observable time [18, 19, 21, 68–70], and its size fluctuates about the carrying capacity, with 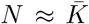. In this setting, it is reasonable to assume 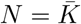 (Moran approximation); see Eqs. (10). The dynamics of the population make-up is thus described by tracking the number *N*_*S*_ of *S* cells present in a population of size 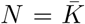 evolving according to the Moran process [8, 53]. (The number of *F* is 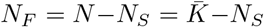.) Within this Moran approximation, the population makeup 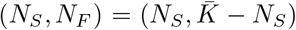 changes according to [8, 53–56]

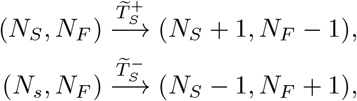

where the Moran transition rates 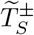 are defined in terms of those of Eq. (2) with 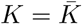 [18–21, 24–28]:

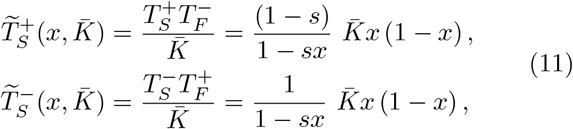

With 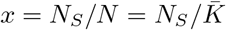. These Moran transition rates correspond to the increase and decrease of the number of *S* by a unit.

Since the population size is constant, every birth of an *S* is accompanied by the death of an *F*, and vice versa. This Moran process is characterised by the absorbing states 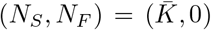 (*S* fixation) and 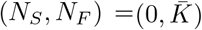 (*F* fixation). The probability and mean time of fixation of this process can be computed analytically [8, 54–56]. When the initial fraction of *S* individuals is 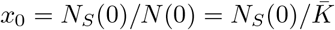, the fixation probability of *S* in a finite population of size 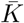 is [21, 25, 27, 55, 56]

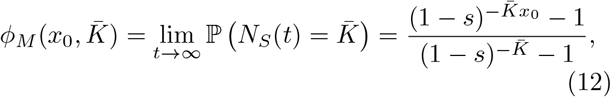

where ℙ (·) denotes the probability that the condition (·) is satisfied. The unconditional Moran mean fixation time, denoted by 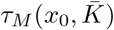, can also be computed exactly [21, 25, 27, 55, 56, 71] and its expression is given of Eq. (B1) in Appendix B.

In what follows, *x*_0_ will be treated as a parameter, and we simplify the notation by generally writing respectively 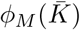 and 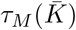 for 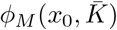 and 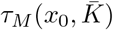, with the value of *x*_0_ clear from the context.

## III. RESULTS: POPULATION SIZE DISTRIBUTION AND FIXATION IN FLUCTUATING ENVIRONMENTS

In the mean-field setting, the slow strain *S* unavoidably goes extinct by Gause’s competitive exclusion principle [72, 73]; see Eq. (10). When *K* is constant and finite, *S* has a fixation probability that vanishes exponentially in a static environment; see Eq. (12). This picture radically changes in fluctuating environments where the carrying capacity varies endlessly by switching back and forth between the values 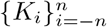 [18–28].

Here, to understand how the environmental variability statistics influence the dynamics of the multi-state switching models, we study the impact of 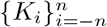 and switching rates 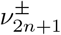 on the population size distribution (PSD), *S* fixation probability and unconditional mean fixation time. According to Eq. (6), the distribution of 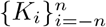, and therefore the amplitude of EF, is controlled by *K*_0_, *K*^±^ and *n*. For the comparison with the effective binary environments, where the carrying capacity switches at effective rates 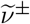 between the extreme values *K*^±^ according to 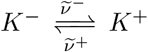, the relation between 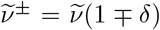 and the multi-state switching rate *ν*_2*n*+1_ is set by Eq. (8).

Our simulation data are obtained by sampling 10^5^ realizations and considering 50-100 values of *ν*_2*n*+1_ in the ranges [10^−2^, 10^2^], [10^−3^, 10], and [10^−4^, 10]. In Figs. 2-8, (*K*^−^, *K*^+^) ∈ {(50, 450), (25, 475)}, *δ* ∈ {0, ±0.1, ±0.2, ±0.25}, *ϵ* ∈ {−0.8, −0.5, −0, 0.5, 5} and (*s, x*_0_) ∈ {(0.02, 0.5), (0.05, 0.6), (0.1, 0.7)}. Simulation methods and further details are given in Appendix C.

**FIG. 2.**
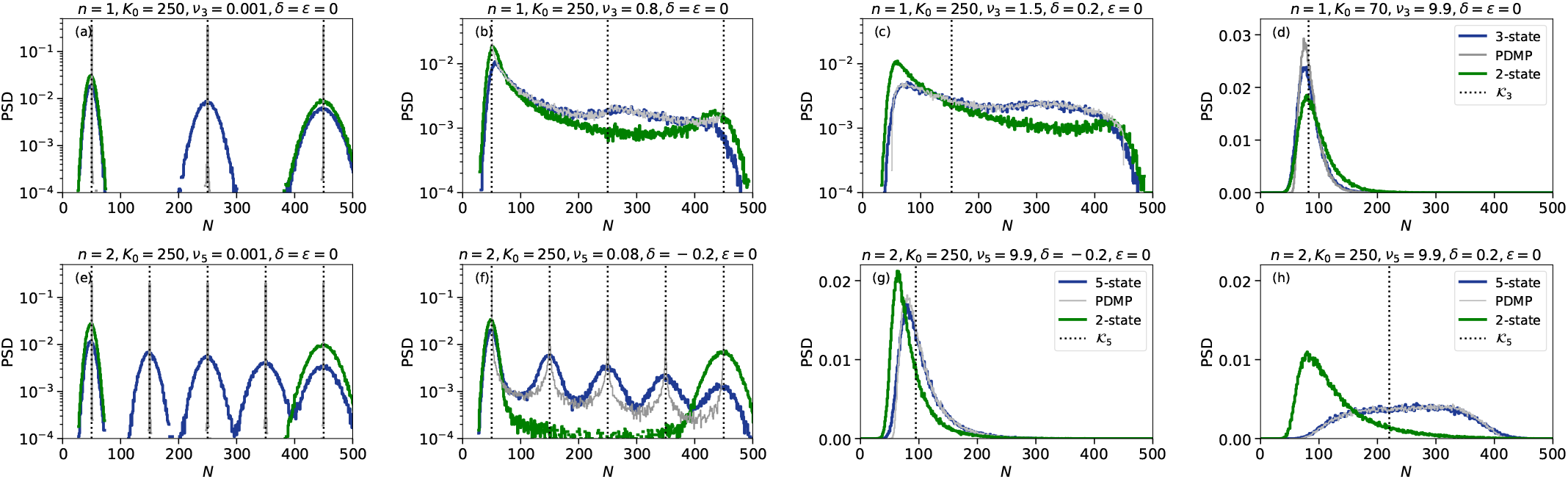
Histograms of the quasi-stationary PSD, 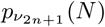 (dark blue), for different switching rates for 3-state (a)(d) and 5-state (b)-(h) multi-state switching models. In grey are shown the histograms of the *N*-PDMP (see Eq, (14) and Appendix C). These are compared with those of the PSD 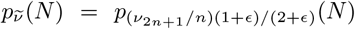 of the effective twostate model (see text and Eq. (8)) shown in green. Panels (a)-(c), (e) and (f) are in semi-log scale (values < 10^−4^ are not shown), while (d), (g) and (h) are in linear scale. The vertical dotted lines in (a),(b),(d),(e) are eyeguides showing *N* = *K*_*i*_. In panels (c) (d),(g),(h) the dotted lines show *K*_3_ ≈ 153.23 (c) and *K*_3_ ≈ 82.17 (d), while in (g) and (h) they respectively indicate *K*_5_ ≈ 94.70 and *K*_5_ ≈ 220.18. In all panels, (*s, x*_0_, *K*^−^, *K*^+^) = (0.02, 0.5, 50, 450). For the green histograms in (d),(g),(h), *K*_2_ = 90 (d), *K*_2_ ≈ 77.59 (g), and *K*_2_ ≈ 107.14 (h); see Eq. (A3). Other parameters are: (*ν*_3_, *K*_0_, *δ, ϵ*) = (10^−3^, 250, 0, 0) in (a), (0.8, 250, 0, 0) in (b), (1.5, 250, 0.2, 0) in (c), (9.9, 70, 0, 0) in (d); and (*ν*_5_, *K*_0_, *δ, ϵ*) = {(10^−3^, 250, 0, 0) in (e), (0.08, 250, −0.2, 0) in (f), (9.9, 250, −0.2, 0) in (g), (9.9, 250, 0.2, 0)} in (h). In (a) and (c) the dotted lines coincide with *N*-PDMP results (grey lines). All histograms have been obtained after *t >* 2000, as outlined in Appendix C. See also Figs. 9(a),(b).

### A. Population size distribution in fluctuating environments

This class of models is characterised by a long-lived, or quasi-stationary, population size distribution (PSD) followed by the eventual extinction of the entire population after a very long time (practically unobservable when *K* ≫ 1 [18, 19, 21, 68]). Here, we focus on timescale *t* ≳ 1*/s* on which one strain is likely to have taken over the population (fixation) while the other has gone extinct, and the population has settled into its long-lived PSD [18–28]; see Fig. 1(b) and Appendix B.

In fluctuating environments, the endlessly switching carrying capacity drives the population size, with *N* varying according to a birth-death logistic process 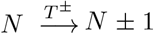, with transition rates 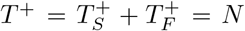 and 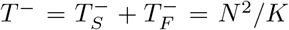. This process has an absorbing boundary at *N* = 0 leading to the eventual collapse of the population after a time that grows dramatically with the system size [18, 19, 21, 68]. Here, have *K* ≥ 25 which is sufficiently large to ensure that the population will not go extinct in our simulations. Here, we first focus the (marginal) PSD, denoted by 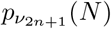, with *δ* and *ϵ* treated as parameters; see Appendices A 3. In Fig. 2, this quantity is compared with the (marginal) PSD of the effective binary switching model, denoted by 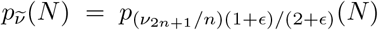 [18, 19, 21, 68]; see Eq. (8) and Appendix A 1. Since *N* relaxes on a timescale *t* ~ 1; see Eq. (10), we distinguish three main regimes [18–21]:

i. When *ν*_2*n*+1_ ≪ 1, environmental switching is much slower than the ecological timescale (“slow switching”). *N* is thus approximately constant and close to one value of 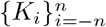 randomly drawn from *π*_*i*_ at time *t* = 0, i.e. *N* ≈ *K*_*i*_ with probability *π*_*i*_. The PSD is therefore multimodal with 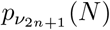 characterised by 2*n* + 1 peaks about each *K*_*i*_, their intensity being set by *π*_*i*_; see Fig. 2(a),(e),(f).
ii. When *ν*_2*n*+1_ ≫ 1, the environment varies on a much faster timescale than the ecological dynamics (“fast switching”). In this regime, *N* is not able to track *K*(*t*) and environmental fluctuations self-average, with the population experiencing the effective carrying capacity *K*_2*n*+1_ whose inverse is obtained by averaging 1*/K* over ***π*** [18–28] (see also Appendix A 1), yielding

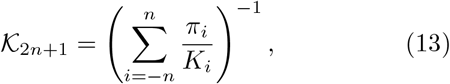

where *K*_*i*_ and *π*_*i*_ are given by Eqs. (6) and (9). The binary counterpart of *K*_2*n*+1_, denoted by *K*_2_, is given by Eq. (A3). In this fast-switching regime, 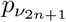 is unimodal and approximately centred at _2*n*+1_; see Fig. 2(c),(g).
iii. When *ν*_2*n*+1_ ~ 1, the time scales of environmental and ecological dynamics are similar (“intermediate switching”) and the PSD has a complex shape; see Fig. 2(b),(c). When *ν*_2*n*+1_ ≲ 1, *N* tracks the time-switching *K*(*t*); see Fig. 1(b), yielding “population bottlenecks” when transitions *K*_*i*+1_ → *K*_*i*_, with *K*_*i*+1_ ≫ 1, occur [24, 27, 28]. Bottlenecks are of great relevance for bacterial dynamics as they lead to new colonies consisting of a small number of individuals prone to fluctuations [24, 27–31]. When *ν*_2*n*+1_ ≳ 1, the peaks of 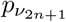 gradually merge; see Fig. 2(c), and as 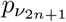 increases the PSD morphs from being multimodal to unimodal.

The blue histograms of Fig. 2 illustrate how environmental and demographic noise jointly affect the PSD in multi-state feast–famine cycles. Clearly, due to random fluctuations, the population size can be above *K*^+^ and below *K*^−^, and thus *p*_*ν*_ is not bounded to [*K*^−^, *K*^+^]. Under slow switching, *ν*_2*n*+1_ ≪ 1, the intensity of the 2*n* + 1 peaks of 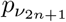, set by *π*_*i*_, is higher for low *K*_*i*_ when *δ* < 0 (Fig.2(f)) and for high *K*_*i*_ when *δ* > 0. The PSD is right-tailed and the peaks around low *K*_*i*_ are always sharper and narrower than those about higher *K*_*i*_; see Fig.2 (a),(b),(c),(e),(f). This stems from the fast decay and slower growth of *N* that is typical of logistic dynamics [18]. In the intermediate switching regime, *ν*_2*n*+1_ ~ 1, the PSD peaks gradually merge, while 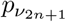 remains right-tailed and has an approximately flat profile between *N* ≈ *K*^±^; see Fig.2(b),(c). In the fast switching regime, *ν*_2*n*+1_ ≫ 1, when *δ* ≤ 0, 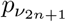 is a righttailed distribution sharply centred about *K*_2*n*+1_ given by Eq. (13); see Fig.2(d),(g). When *δ* > 0 and *n* > 1, the mild environmental states are likely to be populated, resulting in a broader PSD and the convergence towards *N* ≈ *K*_2*n*+1_ requires a very high switching rate; see Fig.2(h). Although *ϵ* modifies the environmental switching dynamics (Appendix A 2), it leaves the stationary distribution ***π*** unchanged; see Eq. (9). Stochastic simulations show no noticeable dependence of the PSD on *ϵ* (for −0.8 ≤ *ϵ* ≤ 5), as illustrated by the comparison of Figs. 2(g),(h) and 9(a),(b), suggesting that 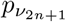 and related quantities are chiefly controlled by the stationary environmental statistics.

The comparison of the multi-state and two-state switching models PSD in Fig. 2 shows that the most striking differences arise in the slow switching regime, where, due to the intermediate environmental states, 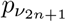 has 2*n* − 1 peaks at *N* ≈ *K*_1−*n*_, …, *K*_*n*−1_ that are absent from the binary PSD; see Fig. 2(a),(e),(f). Moreover, when *ν*_2*n*+1_ ~ 1, the probability that *N* takes a value between *K*^−^ and *K*^+^ is higher in multi-state than binary environments; see Fig. 2(b),(c). Under fast switching, the multi-state and binary-state PSD are both unimodal, but are respectively centred at *K*_2*n*+1_ and *K*_2_, given by Eqs. (13) and (A3), generally with *K*_2*n*+1_ > *K*_2_ when *δ* > 0 (see Figs. 2(h) and 9(b); and see below).

Many features of the quasi-stationary PSD are captured by the piecewise deterministic Markov process [18, 19, 43, 74] for the population size (*N*-PDMP). This process is obtained by letting *N* evolve deterministically between two environmental switches, according to the logistic equation with carrying capacity *K* = *K*_*i*_ in the environmental state *ξ*(*t*) = ∈ *i* {− *n*, …, *n*} until an environmental switch occurs [18–21, 23–28]; see Appendix A 1. Hence, by generalizing the binary *N*-PDMP process of Eq. (A2), the (2*n* + 1)-state *N*-PDMP is defined by

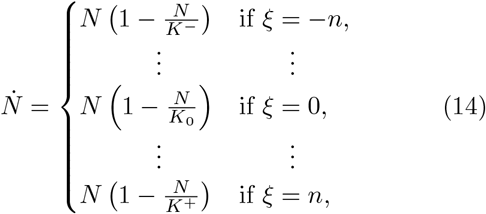

with 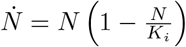 when *ξ*(*t*) = *i* ∈ {−*n*, …, *n*}. The marginal stationary probability density of the binary *N*PDMP (A2) can be computed explicitly and is given by Eq. (A4). For (2*n* + 1)-state switching models, the *N*-PDMP marginal stationary probability density, denoted by 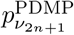, cannot be obtained analytically but can be efficiently computed from simulations of Eq. (14); see Appendix C and Fig. 2.

Since the *N*-PDMP ignores demographic fluctuations, it accounts for randomness only stemming from the random switching of *K*(*t*) and the support of 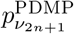 is *N* ∈ [*K*^−^, *K*^+^]. Figure 2 illustrates that 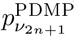 correctly reproduces all the peaks of 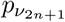; see Fig. 2(a),(b), while it cannot capture the width of the distribution under slow switching. Under intermediate switching, systematic deviations between 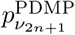 and the PSD mainly arise around *N* ≈ *K*^±^ and are caused by demographic fluctuations; see Fig. 2(b). When *ν*_2*n*+1_ ≳ 1, we find that 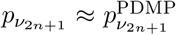 indicating that most of the fluctuations arise from environmental switching and are captured by the *N*-PDMP; see Figs. 2(c), (d),(g),(h) and Fig. 9. The PDMP stationary density also provides an accurate approximation of the average long-term population size 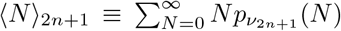, that can be computed numerically as

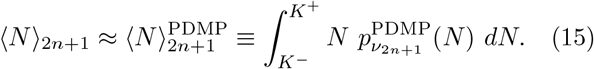

In Figure 3, we find an excellent agreement between ⟨*N*⟩_2*n*+1_ computed from stochastic simulations data and the numerical computation of 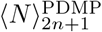 (outlined in Appendix C). As in the binary switching case [18–21, 25, 27], the *N*-PDMP approximation captures that 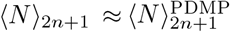 is a decreasing function of *ν*_2*n*+1_ (at fixed *δ, ϵ*), with ⟨*N*⟩_2*n*+1_ ≈ ⟨*K*⟩ _2*n*+1_ when *ν*_2*n*+1_ ≪ 1 and ⟨*N*⟩ *K*_2*n*+1_ ≈ *K*_2*n*+1_ when *ν*_2*n*+1_ ≫ 1. At fixed *ν*_2*n*+1_, ⟨*N*⟩_2*n*+1_ increases with *δ*. In Fig. 3, we notice that the convergence *N* _2*n*+1_ ≈ *K*_2*n*+1_ is slower for *n* = 2 than for *n* = 1 when (*δ, ϵ, K*_0_) are kept fixed; see Fig. 3(a),(c) and Fig. 3(b),(d). This stems from the number of the number of intermediate states increasing with *n* and the effective rate 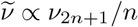; see Eq. (8).

**FIG. 3.**
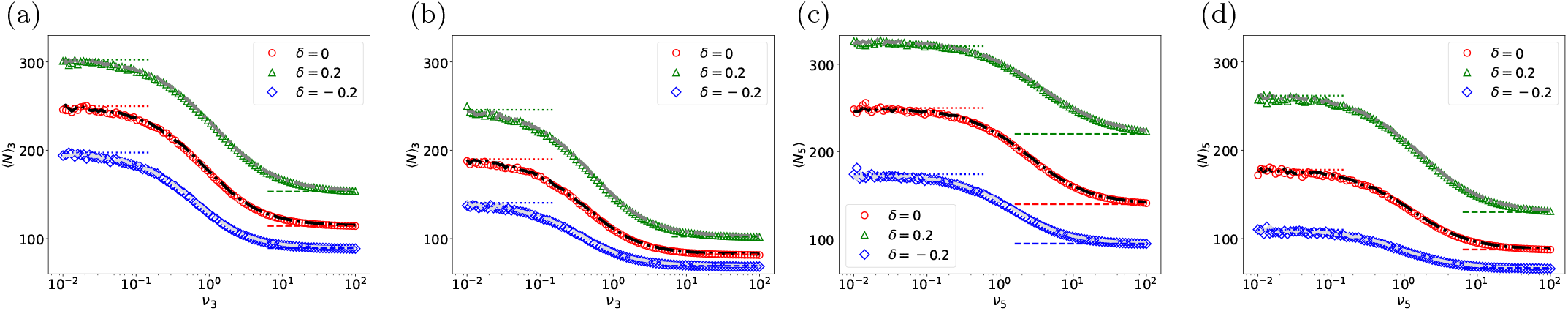
Average population size ⟨*N*⟩ _2*n*+1_ (markers) and 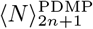 (dash-dotted curves) versus *ν*_2*n*+1_ for different values of *n, K*_0_ and *δ*; see text and (15): (a) (*n, K*_0_) = (1, 250); (b) (*n, K*_0_) = (1, 70); (c) (*n, K*_0_) = (2, 250); (d) (*n, K*_0_) = (2, 70). In all panels, we have *δ* = 0 (red circles, black curves), *δ* = 0.2 (green triangles, dark gray curves), *δ* =−0.2 (blue diamonds, grey curves). The dotted and dashed horizontal lines are eyeguides respectively showing 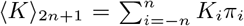 and *K*_2*n*+1_ (given by Eq. (13)) for each combination of *n, K*_0_, *δ*. Other parameters are (*K*^−^, *K*^+^, *ϵ*) = (50, 450, 0) in all panels. In all the examples, ⟨*N* ⟩_2*n*+1_ and 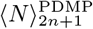 have been computed after *t >* 5000, as outlined in Appendix C.

### B. Fixation in fluctuating environments

In this section, we study how environmental statistics, encoded in 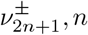, and 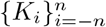, influence the *S* strain fixation probability, denoted by *ϕ*_2*n*+1_ = ℙ (*N*_*S*_(*t*) = *N*(*t*) > 0| *t* < ∞). This is the probability that *S* takes over the entire population in a finite time, avoiding the risk of overall extinction. In Appendix B, we investigate the effect of the environmental statistics on the unconditional mean fixation time (uMFT) for either of the two strains to take over the population.

#### 1. Fixation under slow and fast environmental switching

When *ν*_2*n*+1_ ≪ *s* (slow switching), environmental switching is much slower than selection dynamics. It is therefore is likely that no switches occur on the evolutionary timescale *t* ~ 1*/s* (see Eq. (10) and Appendix B). In this regime, the population is thus subject to the carrying capacity *K*_*i*_ with a probability *π*_*i*_, and an approximate expression of *ϕ*_2*n*+1_ is obtained from the Moran expression (12) by writing 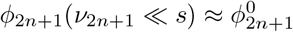, where

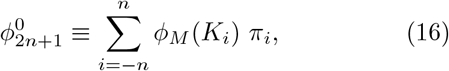

*K*_*i*_ and *π*_*i*_ being given by Eqs. (6) and (9). This corresponds to a “quenched approximation” [26, 75, 76], assuming that fixation occurs while *N* ≈ *K*_*i*_ with a probability *π*_*i*_. A similar reasoning for the binary switching model yields 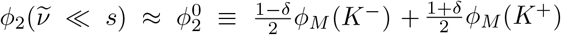 [18, 19, 21, 23–28]; see Appendix A 1.

When *ν*_2*n*+1_ ≫ *s* (fast switching), there are many environmental switches on the timescale *t* ~ 1*/s* yielding the self-average of environmental fluctuations. The carrying capacity hence takes the effective value *K*_2*n*+1_ given by Eq. (13) and *N* ≈ *K*_2*n*+1_. In the fast switching regime, we can thus use an “annealed approximation” [26, 75, 76] for the *S* fixation probability by writing in terms of the Moran result (12) 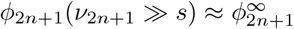, with

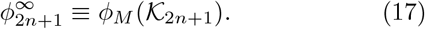

Similarly, in the binary environment we have 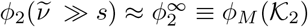 [18, 19, 21, 23–28], where *K*_2_ is given by Eq. (A3).

The heatmaps of Fig. 4(a)-(c) and Fig. 4(d)-(f) respectively show the ratios 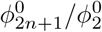 and 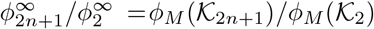 for different values of *δ* as a functions of *n* and *K*_0_ (for *s, x*_0_ and *K*^±^ kept fixed). The regions where 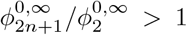 (orange/pink) are separated from those where 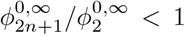 (blueish) by dashed lines along which 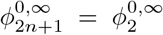. Due to the strong dependence of *π*_*i*_ on *δ*; see Eq. (9), the shape and area of these regions vary greatly with *δ*. When *δ* ≤ 0 and *K*_0_ is sufficiently low, the *S* fixation probability under slow and fast switching can be higher in multi-state than in binary environments for all *n* ≥ 1; see Fig. 4(a),(c),(d),(f). When *δ* < 0, environmental bias is towards states having a low carrying capacity, resulting in predominantly orange/pink-dominated heatmaps where 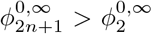 with 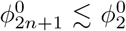 for sufficiently high values of *K*_0_ (pale blue below the dashed line); see Fig 4(c,f). When *δ* > 0, environmental switching is biased towards states with a high carrying capacity, yielding the predominantly blue-dominated heatmap of Fig. 4(b,d), corresponding to 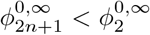 for most o*n, K*_0_ pairs. (For example, 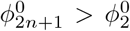 only for *K*_0_ ≲ 100 when *n* ≥ 2). The results of Fig. 4 are in agreement with the simulation data of Figs. 5 and 6, and illustrate that the distribution of carrying capacities 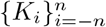 (here set by *n* and *K*_0_, *K*^±^) can lead to complex fixation scenarios, significantly richer than those arising under binary switching.

**FIG. 4.**
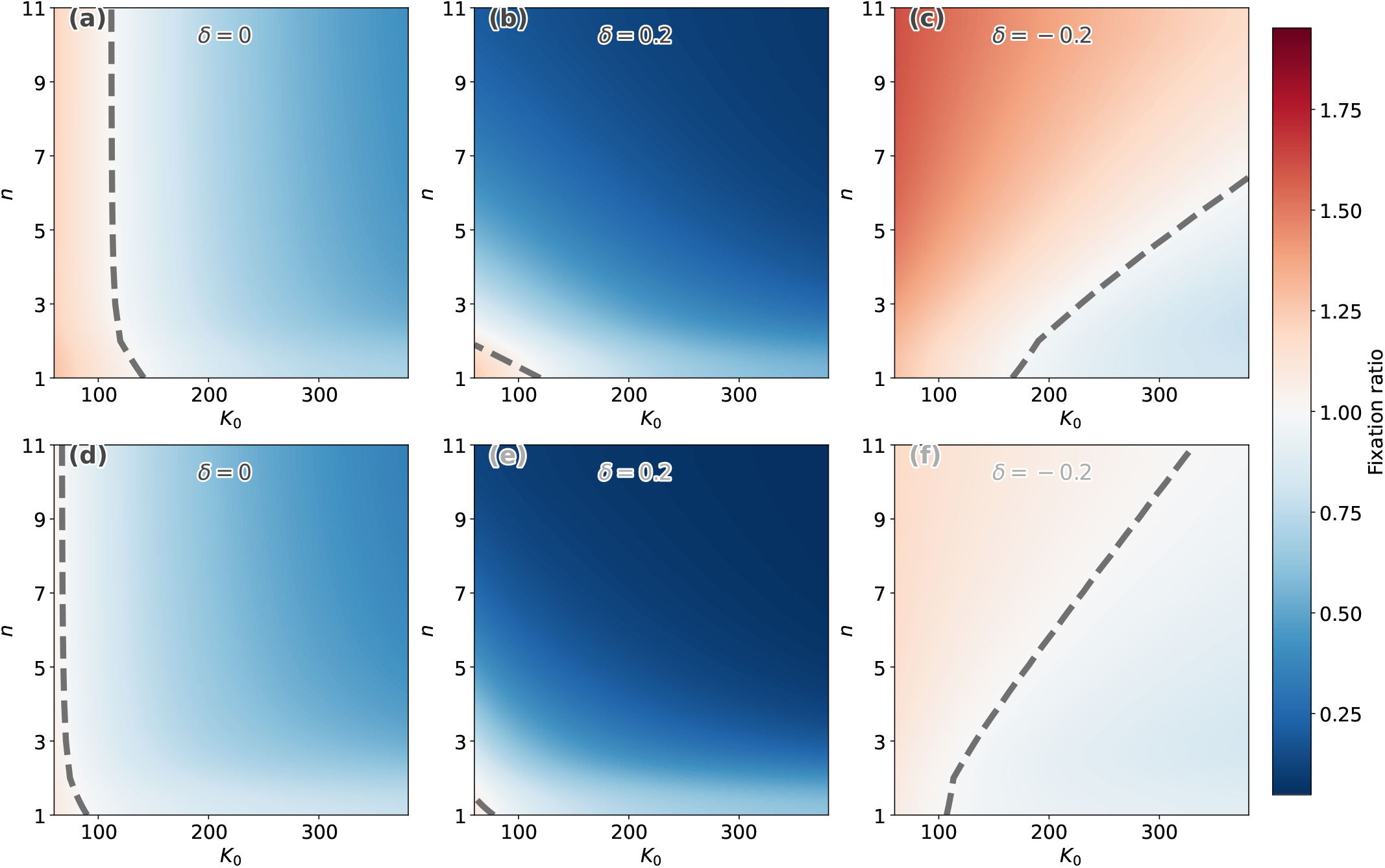
Comparison of the *S* fixation probability in the regime of slow and fast switching in multi-state (*n* ≥ 1) and binary switching models: Heatmaps of the ratios 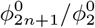 (slow switching) in (a)-(c) and 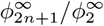 (fast switching) in (d)-(f) as colour-coded in the rightmost vertical bar; see text. Parameters are (*K*^−^, *K*^+^, *s, x*_0_, *ϵ*) = (50, 450, 0.02, 0.5, 0) and *δ* = 0 in (a,d), *δ* = 0.2 in (b,e), and *δ* = −0.2 in (c,f). For reference, *K*_2_ = 90 for *δ* = 0, *K*_2_ ≈ 107.14 for *δ* = 0.2, and *K*_2_ ≈ 77.59 for *δ* = −0.2. In all panels, 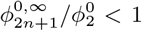 blueish areas and 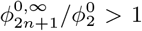 in yellow/orange regions. The black dashed lines (surrounded by a whitish cloud) indicate the contours along which the ratios equal to one.

**FIG. 5.**
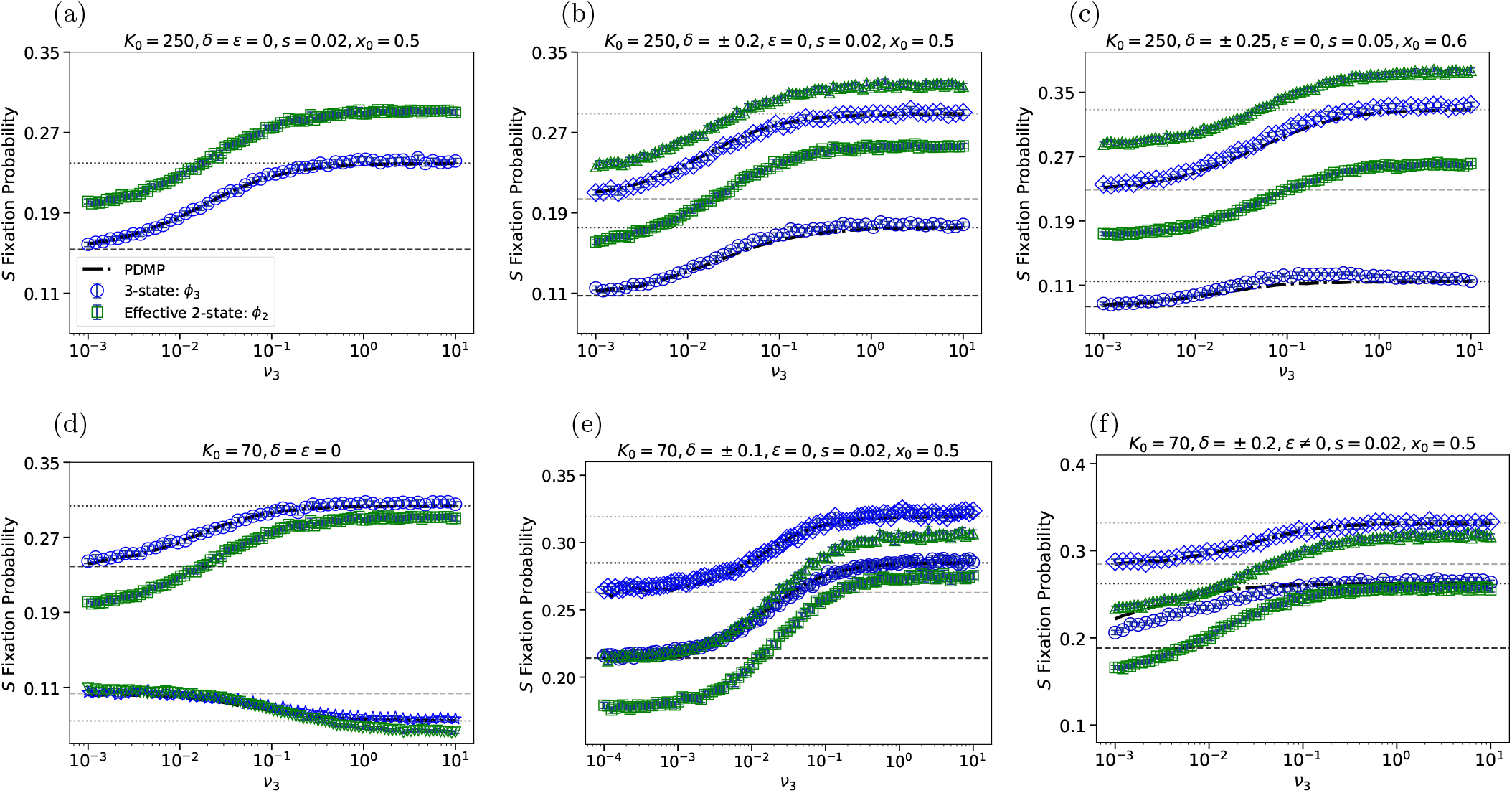
*S* fixation probability in the three-state model as a function of the switching rate *ν*_3_. The centre state carrying capacity in (a)-(c) is *K*_0_ = 250, while *K*_0_ = 70 in (d)-(f). Here, *n* = 1, and (*K*^−^, *K*^+^) = (50, 450) in all panels except (c). Symbols are from full stochastic simulations and black dashed-dotted curves (in all panels), often almost indistinguishable from markers, are from the *N*-PDMP-based approximation; see Eq. (19). In blue, is shown *ϕ*_3_ vs. *ν*_3_ for different parameter sets: (*δ, ϵ, s, x*_0_) = (0, 0, 0.02, 0.5) (circles) in (a); (*δ, ϵ, s, x*_0_) = (−0.2, 0, 0.02, 0.5) (circles) and (*δ, ϵ, s, x*_0_) = ( 0.2, 0, 0.02, 0.5) (diamonds) in (b); (*δ, ϵ, s, x*_0_, *K*^−^, *K*^+^) = (0.25, 0, 0.05, 0.6, 25, 475) (circles) and (*δ, ϵ, s, x*_0_, *K*^−^, *K*^+^) = (− 0.25, 0, 0.05, 0.6, 25, 475) (diamonds) in (c); (*δ, ϵ, s, x*_0_) = (0, 0, 0.02, 0.5) (circles) and (*δ, ϵ, s, x*_0_) = (0, 0, 0.1, 0.7) (crosses) in (d); (*δ, ϵ, s, x*_0_) = (0.1, 0, 0.02, 0.5) (circles) and (*δ, ϵ, s, x*_0_) = (−0.1, 0, 0.02, 0.5) (diamonds) in (e); (*δ, ϵ, s, x*_0_) = (0.2, 5, 0.02, 0.5) (circles) and (*δ, ϵ, s, x*_0_) = (−0.2, 0.5, 0.02, 0.5) (diamonds) in (f). Green markers show simulation data for the fixation probability *ϕ*_2_ of the effective two-state model; see text and Eq. (8). In (b,c,e,f), results for *ϕ*_2_ are shown as triangles when *δ* = −0.2 and as squares when *δ* = 0.2. In (d), simulations data of *ϕ*_2_ are shown as squares for (*s, x*_0_) = (0.02, 0.5) and as downside triangles for (*s, x*_0_) = (0.1, 0.7). For the *N*-PDMP-based approximation of *ϕ*_2_, see Refs. [18, 19, 21]. Horizontal dashed and dotted lines are eyeguides showing 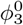 (dashed) and 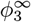 (dotted) from Eqs. (16) and (17) for each parameter set. Error bars are included in each case but are typically too small to see (Appendix C).

**FIG. 6.**
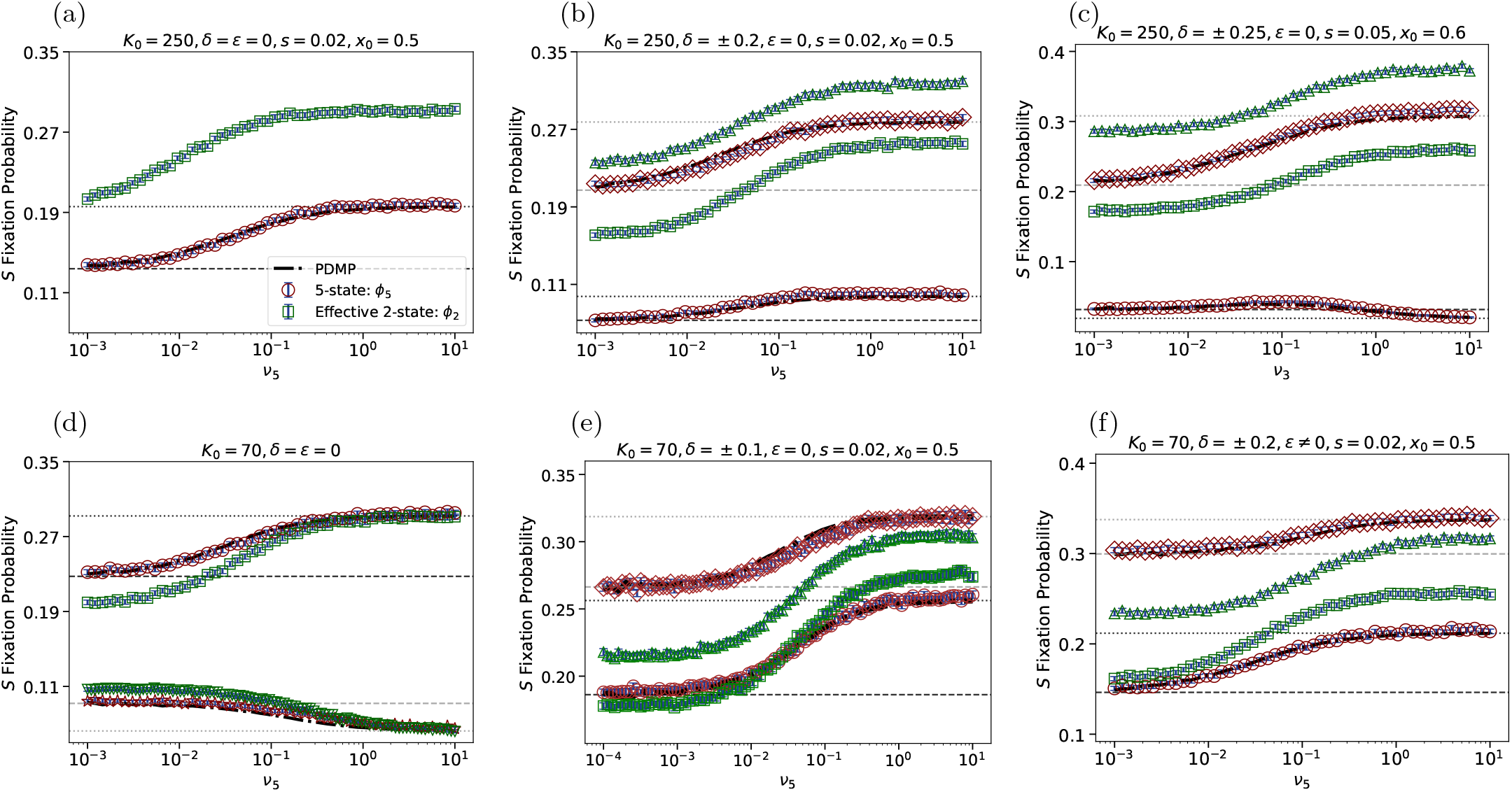
*S* fixation probability in the five-state model as a function of the switching rate *ν*_5_. The centre state carrying capacity in (a)-(c) is *K*_0_ = 250, while *K*_0_ = 70 in (d)-(f). Here, *n* = 2, and (*K*^−^, *K*^+^) = (50, 450) in all panels except (c). Symbols are from full stochastic simulations and black dashed-dotted curves (in all panels), almost indistinguishable from markers, are from the *N*-PDMP-based approximation; see Eq. (19). In brown, is shown *ϕ*_5_ vs. *ν*_5_ for different parameter sets: (*δ, ϵ, s, x*_0_) = (0, 0, 0.02, 0.5) (circles) in (a); (*δ, ϵ, s, x*_0_) = (0.2, 0, 0.02, 0.5) (circles) and (*δ, ϵ, s, x*_0_) = (−0.2, 0, 0.02, 0.5) (diamonds) in (b); (*δ, ϵ, s, x*_0_, *K*^−^, *K*^+^) = (0.25, 0, 0.05, 0.6, 25, 475) (circles) and (*δ, ϵ, s, x*_0_, *K*^−^, *K*^+^) = ( 0.25, 0, 0.05, 0.6, 25, 475) (diamonds) in (c); (*δ, ϵ, s, x*_0_) = (0, 0, 0.02, 0.5) (circles) and (*δ, ϵ, s, x*_0_) = (0, 0, 0.1, 0.7) (brown crosses) in (d); (*δ, ϵ, s, x*_0_) = (0.1, 0, 0.02, 0.5) (circles) and (*δ, ϵ, s, x*_0_) = (−0.1, 0, 0.02, 0.5) (diamonds) in (e); (*δ, ϵ, s, x*_0_) = (0.2, 5, 0.02, 0.5) (circles) and (*δ, ϵ, s, x*_0_) = (− 0.2, −0.5, 0.02, 0.5) (diamonds) in (f). As in Fig. 5, green markers show simulation data for the fixation probability *ϕ*_2_ of the effective two-state model; see text and (8). In (b,c,e,f), results for *ϕ*_2_ are shown as triangles when *δ* = 0.2 and as squares when *δ* = −0.2. In (d), simulations data of *ϕ*_2_ are shown as squares for (*s, x*_0_) = (0.02, 0.5) and as downside triangles for (*s, x*_0_) = (0.1, 0.7). For the *N*-PDMP-based approximation of *ϕ*_2_, see Refs. [18, 19, 21]. In (c), *ϕ*_5_ varies weakly but non-monotonically with *ν*_5_. Horizontal dashed and dotted lines are eyeguides showing 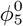 (dashed) and 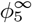 (dotted) from Eqs. (16) and (17) for each parameter set. Error bars are included in each case but are too small to see (Appendix C).

In binary switching environments, the Moran approximations 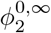 were found to work better under weak selection, *s* ≪ 1, because fixation then occurs on a slower timescale [19]. The evolutionary dynamics of *x*(*t*) therefore samples many demographic fluctuations of *N*(*t*), so that deviations from an effectively constant population size approximately average out. For stronger selection, *s* = *O*(1), fixation events occur more rapidly and the coupling between fluctuations of *N*(*t*) and *x*(*t*) leads to larger deviations from the Moran predictions. As a result, the slow/fast-switching approximations 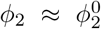 and 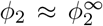 are more accurate when *s* ≪ 1 than for *s* = *O*(1) [19]. In contrast, for multi-state environments, switches occur through intermediate carrying capacities.

This gradual variation reduces the influence the selection strength, yielding *ϕ*_2*n*+1_ to be well approximated by 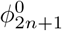 and 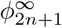 for weak and moderate selection strengths in the suitable switching regimes; see Figs. 5(d) and 6(d).

#### 2. Fixation under intermediate environmental switching

Under intermediate environmental switching, where *ν*_2*n*+1_ ~ *s* (|*δ*| < 1), evolutionary and environmental dynamics take place on similar time scales, and *N* has a nontrivial PSD; see 2(b),(c). It is generally difficult to obtain analytical results for the fixation in the intermediate switching regime. Here, in addition to stochastic simulations, we devise an efficient *N*-PDMP approximation, generalizing the binary-state approach of Refs. [18– 20] (see Appendix A 1). The method relies on a timescale separation, and a suitable rescaling of the switching rate. When *s* ≪ 1, *N* settles in its (quasi-)stationary PSD on a much shorter timescale than *t* ~ 1*/s* (evolutionary timescale); see Eq. (10). *N* is thus considered at quasistationarity, with the *N*-PDMP providing a suitable approximation of its dynamics. In Appendix A 2, we show that the average number of switches on the timescale 1*/s* is 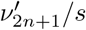, where

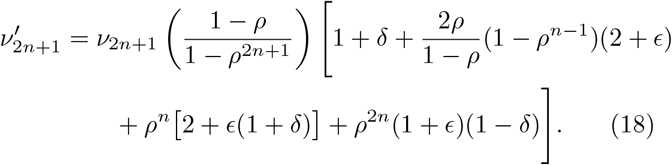

When *δ* = *ϵ* = 0, we have 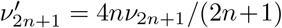. Following Refs. [18–21, 27], we can use the *N*-PDMP density to approximate the *S* fixation probability when *ν*_2*n*+1_ ~ *s* and 1*/* ⟨*N*⟩ ≪ *s* ≪ 1. For this we average the Moran fixation probability *ϕ*_*M*_ (*N*), obtained from Eq. (12), over 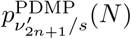, where the switching rate has been rescaled, 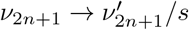 using Eq. (18), to properly account for the expected number of switches occurring on the fixation timescale [18–21, 23, 27].:

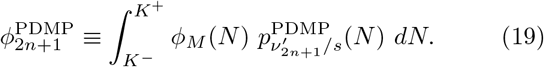

In Appendix B, this approach is used for the uMFT. A similar approximation for *ϕ*_2_ and uMFT in binary environments has been studied in Refs. [18, 19, 21]; see Appendix A 1. In practice, 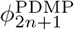 is efficiently computed numerically as outlined in Appendix C. The comparison with extensive simulations of the individual-based models; see Figs. 5 and 6, shows that Eq. (19) generally gives a very good approximation of the *S* fixation probability, with 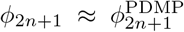 over a broad range of values of the switching rate, *ν*_2*n*+1_ ≪~ *s*, as well as *ν*_2*n*+1_ ≪ *s* and *ν*_2*n*+1_ ≫ *s* (slow/fast switching regimes). The same agreement is found in binary-switching environments [18–20, 27].

Below, for the sake of concreteness, we discuss in some detail the cases of ternary and five-fold fluctuating environments (*n* = 1 and *n* = 2).

#### 3. Fixation in the three-state switching model

When *n* = 1, the ternary environmental cycle consists of the feast and famine states of carrying capacity *K*^±^, and the median state whose carrying capacity is *K*_0_. Thus, in the three-state switching model *K*(*t*) ∈ {*K*^−^, *K*_0_, *K*^+^}; see Fig. 1(a,left).

At the start of each simulation, the value of *K* is randomly allocated according to the distribution given by Eq. (9) which, for *i* ∈ {−1, 0, 1}, explicitly reads

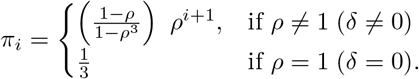

In the regime of slow switching where *ν*_2*n*+1_ ≪ *s*, the *S* fixation probability is approximately given by Eq. (16), with 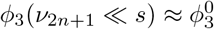, where

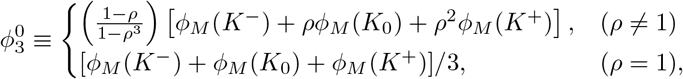

and *ϕ*_*M*_ (*K*_*i*_) is given by Eq. (12). In binary environments, this approximation yields 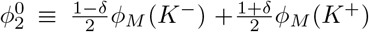 [18–21, 23] (Appendix A 1). From the comparison of these results, we find that the *S* fixation probability is higher in ternary environments 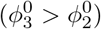 when 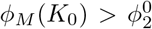, and 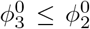 otherwise. The conditions for 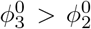 in the examples of Fig. 4(a)-(c) are therefore *K*_0_ ≲ 141.36 (*δ* = 0), *K*_0_ ≲ 119.61 (*δ* = 0.2), and *K*_0_ ≲ 166.50 (*δ* = −0.2). The horizontal dashed lines in Fig. 5 show that 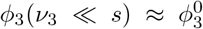, and hence 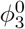 is a good approximation in the slow switching regime, with 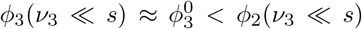 when *K*_0_ = 250 (Fig. 5(a)-(c)), and 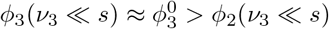 when *K*_0_ = 70 in Fig. 5(e),(f) and Fig. 5(d) when *s* = 0.02.

In the fast switching regime, the population is subject to the effective carrying capacity given by Eq. (13), that here reads

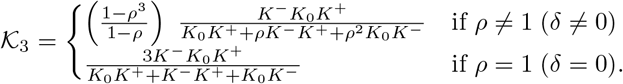

Comparing this expression with its binary-state counterpart, *K*_2_, given by Eq. (A3), we find that *K*_2_ > *K*_3_ when 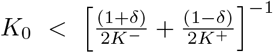. Since *ϕ*_*M*_(*K*) is a decreasing function of *K*, the *S* fixation probability under fast switching is higher in 3-state than 2-state environments 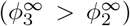 when 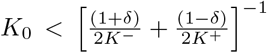, whereas it is the opposite 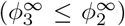 otherwise. This is captured by the heatmaps of Fig. 4(d)-(f) where for *n* = 1 we find 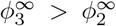 when *K*_0_ < 90 in (d), if *K*_0_ ≲ 77.59 in (e), and when *K*_0_ ≲ 107.14 in (f). The dotted lines in Fig. 5(a)-(c) illustrate that, for given (*K*^±^, *s, x*_0_, *δ, ϵ*), 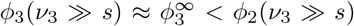 when *K*_0_ = 250; while those in Fig. 5(d)-(f) show that 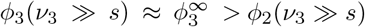 when *K*_0_ = 70.

In the intermediate switching regime (*ν*_3_ ~ *s*), simulation results of Fig. 5 show that the *N*-PDMP-based approximation given by Eq. (19) generally faithfully captures the properties of *ϕ*_3_ in all regimes, with 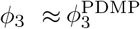. In Fig. 5(a)-(e), dashed lines showing 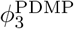 are almost indistinguishable from *ϕ*_3_ for all switching rate values.

Since the environmental switching is biased towards higher population size when *δ* > 0 (Fig. 2) and *ϕ*_*M*_ (*N*) is a decreasing function of *N, ϕ*_3_ decreases with *δ* at fixed value of *ν*_3_ (Fig. 2(b),(f)) and is higher for lower values of *K*_0_; compare Fig. 5(a),(d) and Fig. 5(b),(f). This feature holds for arbitrary values of *n*; see below. We have verified that for − 0.8 ≤ *ϵ* ≤ 5 the parameter *ϵ* has no noticeable effect on *ϕ*_3_; e.g., compare Fig. 5(b) and Fig. 10(a); see Appendix D. For high value of *ϵ*, the *N*PDMP-based approximation overestimates the average number of switches prior to fixation, leading to 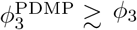 when *ν*_3_ ≲ *s* in Fig. 5(f) and Fig. 10(a) when *ϵ* = 5. These results show that, depending on the value of *K*_0_, ternary switching can either enhance or decrease the *S* fixation probability with respect to the binary case: When *K*_0_ is sufficiently low (e.g. *K*_0_ = 70 in Fig. 5(d)(f), the population size is small over significant periods, during which the selection pressure is balanced by demographic fluctuations. This leads to the *ϕ*_3_ to be larger than *ϕ*_2_ and closer to the fixation probability of the faster strain.

#### 4. Fixation in the five-state switching model

When *n* = 2, the five-fold environmental cycle consists of the feast, famine and median states of respective carrying capacities *K*_±2_ = *K*^±^ and *K*_0_, and the intermediate mild/harsh states of carrying capacity *K*_±1_ = (*K*_0_ + *K*^±^)*/*2. Hence, in the five-state switching model *K*(*t*) = ∈ {*K*_*i*_ *K*^−^, *K*_−1_, *K*_0_, *K*_1_, *K*^+^}; see Fig. 1(a,right), with its value randomly initiated from the distribution given by Eq. (9).

In the regime of slow switching where *ν*_2*n*+1_ ≪ *s*, the *S* fixation probability is approximated by 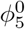, from Eq. (16). For given values of *s, x*_0_, *δ, K*^±^, by solving 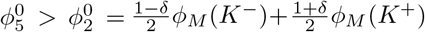, we find the conditions on *K*_0_ for which the *S* fixation probability under slow switching is enhanced by five-state environments with respect to the binary case. For the examples of Fig. 4(a)-(c) with *n* = 2, we find 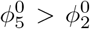 if *K*_0_ ≲ 119.96 (*δ* = 0), *K*_0_ ≲ 52.74 (*δ* = 0.2), and *K*_0_ ≲ 190.22 (*δ* = −0.2).

In the case *n* = 2, the effective carrying capacity under fast switching, given by Eq. (13), explicitly reads 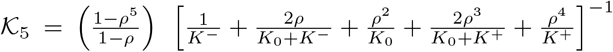 when *ρ ≠* 1 (*δ* ≠ 0), which becomes 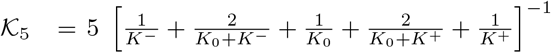 when switching is symmetric (*ρ* = 1, *δ* = 0). Proceeding as in the ternary case, the comparison of *K*_5_ and *K*_2_ leads to the conditions for *K*_2_ > *K*_5_ and thus for 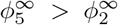. In Fig. 4(d)-(f) we thus find that 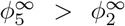 when *K*_0_ < 74.40 (*δ* = 0), if *K*_0_ ≲ 113.34 (*δ* = − 0.2), while 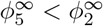 when *δ* = 0.2.

The dotted and dashed lines in Fig. 6 show that 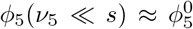 and 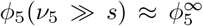, with 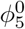 and 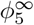 thus providing good approximations in the slow and fast switching regimes when *K*_0_ = 250 and *K*_0_ = 70.

When *ν*_5_ ~ *s* (intermediate switching), Figure 6 shows that the *N*-PDMP-based approximation, given by Eq. (19), is in good agreement with simulation results, yielding 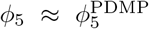 for all parameter sets. Again, we notice that Eq. (19) is a good approximation across the slow, intermediate and fast switching regimes, with dashed-dotted lines in Fig. 6 almost indistinguishable from the data of *ϕ*_5_. In particular, *ϕ*_5_^PDMP^ captures the non-trivial *ν*_5_-dependence of *ϕ*_5_, and even reproduces its weak peak in the intermediate regime (for *ν*_5_ ~ 0.1) of Fig. 6(c). As in ternary environments, *ϕ*_5_ at given *ν*_5_ decreases with *δ*; see Fig. 6(b), and is higher for lower values of *K*_0_; compare Fig. 6(a),(d) for (*s, x*_0_) = (0.02, 0.5) and Fig. 6(b),(f).

The comparison of Figs. 5 and 6, supported by the heatmaps of Figs. 4, indicates that the *S* fixation probability decreases when the number of environmental states increases, with the environmental bias *δ* > 0 strongly reducing *ϕ*_2*n*+1_ as *n* becomes bigger. This effect can be countered by reducing the value of *K*_0_. Generally, *ϕ*_2*n*+1_ and *ϕ*_2_ exhibit the same qualitative dependence on the switching rate, with *n* affecting mostly the amplitude of *ϕ*_2*n*+1_, and *ϕ*_2*n*+1_ < *ϕ*_2_ or *ϕ*_2*n*+1_ > *ϕ*_2_ for all values of *ν*_2*n*+1_. However, for some parameters *ϕ*_2*n*+1_ < *ϕ*_2_ in a certain switching regime and *ϕ*_2*n*+1_ > *ϕ*_2_ in another; see Fig. 6(e) where, for *δ* = 0.1, *ϕ*_5_ > *ϕ*_2_ in the slow/intermediate regime and *ϕ*_5_ < *ϕ*_2_ under intermediate/fast switching. There are also parameter sets for which one of *ϕ*_2*n*+1_ or *ϕ*_2_ varies non-monotonically with *ν*_2*n*+1_ while the other is an increasing or decreasing function of the switching rate (Figs. 5(c) and 6(c)), or where *ϕ*_2*n*+1_ < *ϕ*_2_ and *ϕ*_2*n*+1_ > *ϕ*_2_ on two separate regimes of *ν*_2*n*+1_ (Fig. 6(e)).

These results show that multi-state switching environments lead to various fixation scenarios, with the *S* fixation probability that can be enhanced or hindered with respect to the binary case, depending on the values of *K*_0_ and *ν*_2*n*+1_, encoding the amplitude and frequency of environmental fluctuations.

## IV. DISCUSSION & CONCLUSION

Microbial communities are commonly subject to changing environmental conditions that are often described in terms of two-state switching models. Strain competition subject to a binary time-switching carrying capacity can be seen as the simplest coarse-grained description of feast–famine cycles, characterised by alternating periods of nutrient abundance (feast) and scarcity (famine). Recent experimental and theoretical studies, however, show the importance of *multiple intermediate environmental states* on feast-famine cycle dynamics [48– 50]. Experiments have indeed demonstrated that the frequency and amplitude of environmental fluctuations have a strong impact on microbial feast–famine cycles, with gradual changes of nutrient conditions influencing nutrient-utilization strategies. Eco-evolutionary dynamics in binary and ternary environments revealed important differences between abrupt and gradual environmental deterioration through intermediate conditions [51].

This has motivated us to study a class of multi-state switching models for the dynamics of two strains, one (*S*) slightly slower than the other, competing for the same resources in multi-state fluctuating environments. Here, the time-varying environment consists of multiple ordered (2*n* + 1) environmental states, each associated with its own carrying capacity. To represent gradually changing conditions, there are switches between nearestneighbour states and their carrying capacities: The environment evolves as a Markov chain over multiple carrying capacity states, which provides a simple description of gradual environmental variability. The individual-based switching models considered here are not intended as a mechanistic description of feast–famine experiments, but can be interpreted as a coarse-grained representation of fluctuating nutrient environments. In this picture, the multi-state carrying capacities are associated with different nutrient conditions, the switching rates determine the frequency of environmental fluctuations, and the distribution of carrying capacities quantifies their amplitude. This class of models therefore captures, at the population level, the influence of the statistical properties of environmental variability on eco-evolutionary dynamics, while coarse-graining over microscopic processes such as nutrient uptake. From this perspective, the (2*n*+1)-state switching models provide an individual-based framework for investigating how the frequency and amplitude of environmental fluctuations jointly affect stochastic feast– famine cycle dynamics and their evolutionary outcomes.

By computational and analytical means, we have characterised the long-time dynamics and fate of these multistate switching models. In particular, we have studied the influence of the switching rates and distribution of carrying capacities on the long-time population-size statistics and fixation properties, and compared these results with those obtained in binary environments. Due to the presence of intermediate states, the population size distribution (PSD) is generally broader and the environmental bias (parameter *δ*) has a stronger effect than under binary switching. In general, owed to the underlying logistic dynamics, the PSD is right-tailed. Under slow switching, the PSD is characterised by (2*n* + 1) peaks, whereas fast switching yields the same qualitative behaviour as in two-state environments. Many features of the PSD are aptly captured by a description of the population size by a piecewise-deterministic Markov process (*N*-PDMP). The *S* fixation probability in the multi-state switching models can be either hindered or enhanced with respect to the binary case, depending on the switching rate, number of states and their carrying capacities. Since, the slow strain is more likely to fixate the population under harsh than mild conditions, increasing the number of environmental states and the values of the carrying capacities tends to reduce the *S* fixation probability. However, there are environmental conditions under which the fixation of *S* is more likely in the presence of intermediate environmental states than in the binary case; see Figs. 5(d)-(f) and 6(d)-(f). The unconditional mean fixation time scales with the evolutionary timescale (1*/s*), with the prefactor depending on the frequency and amplitude of environmental variability. The probability and unconditional mean time of fixation can be obtained analytically in the limits of slow and fast switching. We have also devised an *N*-PDMP-based approximation for the fixation properties under weak selection, an efficient computational method that provides useful scaling information.

In the feast–famine cycles considered in Refs. [48–50], environmental variability is encoded in nutrient dynamics. In the class of switching models studied here, multistate carrying capacities provide a coarse-grained description of varying nutrient levels. This individual-based framework has allowed us to analyse how the frequency and amplitude of environmental fluctuations shape population dynamics, highlighting that environmental complexity leads to richer eco-evolutionary scenarios. In the future, this approach could be generalised in different directions. In addition to straightforward extensions (e.g., 2*n* environmental states, other distributions of the carrying capacities), we can mention the limit of a large number of environmental states that will help shed further light on the notoriously difficult problem of ecoevolutionary dynamics under continuous environmental noise; see, e.g., Refs. [3, 23, 45, 77, 78]. This framework can also be employed to investigate the coarse-grained effect of varying nutrient levels on the spread of cooperative antimicrobial resistance in well-mixed settings [24, 79], or in spatially structured metapopulations inspired by chemostat systems or batch cultures [17, 28, 59, 62, 80]. More generally, the present work shows that multi-state stochastic switching environments provide a simple yet versatile framework for investigating how the statistical properties of environmental fluctuations shape ecoevolutionary dynamics.

## Data availability statement

Simulation data and codes that support the findings of this study are openly available at the following URL/DOI: 10.5518/1899 [81].

## ACKNOWLEDGMENTS

Partial support from the U.K. Engineering and Physical Sciences Research Council (EPSRC) under the Grant No. EP/V014439/1 is gratefully acknowledged.

## Appendix A Further details on the class of switching models

### 1. Background: Competition in binary-state switching models

As a background for this study, we review the main features of the competition dynamics in binary-state switching models; see Refs. [18–21] for further details.

In two-state switching models, with cyclically alternating mild and harsh conditions, the time-varying carrying capacity *K*(*t*) ∈ {*K*^−^, *K*^+^} takes only two possible values: *K*^+^ and *K*^−^ representing respectively the carrying capacity of the “feast” and “famine” states (*K*^+^ *> K*^−^) [18–28]. It is thus convenient to write the carrying capacity of two-state switching models as

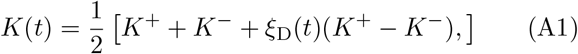

where *ξ*_D_(*t*) ∈ {−1, 1} is the coloured dichotomous Markov noise (DMN) [63–65], also called telegraph process, encoding the binary environmental variability and driving the switching of *K*. The DMN switches between *±*1 according to *ξ*_D_ → − *ξ*_D_ at rate 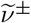 when *ξ*_D_ = *±*1 [63– 65]. It is useful to write 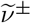 in terms of the mean switching rate 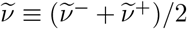 and 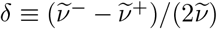 (environmental switching bias), with |*δ*| *<* 1. When the DMN is at stationarity, all instantaneous correlations become time-independent while its autocovariance is a function of the time difference. This means that at stationarity *ξ*_D_ = *±*1 with a probability (1 *± δ*)/2, and the stationary average of the DMN is ⟨*ξ*_D_(*t*) ⟩ = *δ*, while its autocovariance reads 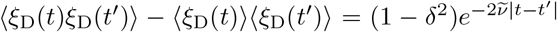 when *t, t*^*′*^ → ∞ [21, 23, 63–65], where ⟨ ⟩ here denotes the ensemble average and 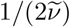 is the finite DMN correlation time.

Following Eq. (A1), the binary carrying capacity switches back and forth at rates 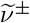 between a value *K* = *K*^+^ (*ξ*_D_ = 1) in a mild environment, and *K* = *K*^−^ *< K*^+^ (*ξ*_D_ = −1) when environmental conditions are harsh, according to 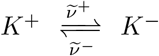 As the DMN, when the time-switching *K*(*t*) is at stationarity, it takes the value *K* = *K*^*±*^ with probability (1 *± δ*)*/*2. Its expected value and variance are time independent, and respectively read 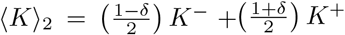, and var(*K*)_2_ = (1 − *δ*^2^)(*K*^*+*^ − *K*^*-*^)^*2*^*/4, while* its auto-covariance is 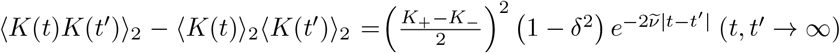 [21, 23, 63–65].

The time-switching *K*(*t*) drives the overall population size *N*(*t*), and is hence responsible for the coupling of demographic and environmental fluctuations [18–28]. Upon ignoring all fluctuations, the mean-field dynamics of the population subject to a constant carrying capacity 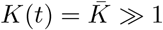 satisfy the rate equations (10) for *N* and the fraction *x* = *N*_*S*_*/N* of *S* individuals; see Sec. II B. When 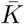 is large but finite, as explained in Sec. II C, the population dynamics can be aptly approximated by a Moran process [8, 12, 13, 18–20, 53, 54] by assuming a constant population size 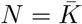. In this case the Moran probability and mean time for the fixation can be computed analytically [18, 19, 21, 24–28, 55, 56]; see Sec. II C and Appendix B 1.

When the population size is sufficiently large for demographic fluctuations to be negligible, but is still subject to environmental variability (switching of *K*(*t*)), the population size dynamics is well approximated by the two-state piecewise deterministic Markov process (*N*-PDMP) [18–21, 23–28, 43, 74] that for binary switching is defined by

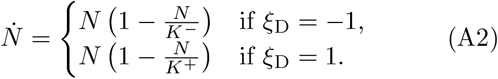

In the *N*-PDMP approximation, the population size satisfies a deterministic logistic equation in each environmental state *ξ*_D_ = *±*1, subject to the time-switching carrying capacity *K*(*t*) given by Eq. (A1). The case of the *N*-PDMP approximation for multi-state switching models with (2*n* + 1) environmental states is discussed in Sec. III A. While it ignores the effect of demographic noise, the *N*-PDMP approximation captures many properties of the (marginal) quasi-stationary distribution 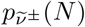 of the population size in binary environments, including that 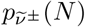 is bimodal under slow switching 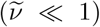, while it is unimodal and centred around the effective carrying capacity

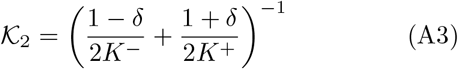

in the fast environmental switching regime 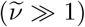 [18– 21, 23–28]. The marginal probability density of the *N*PDMP (A2) can be obtained analytically [18–21]:

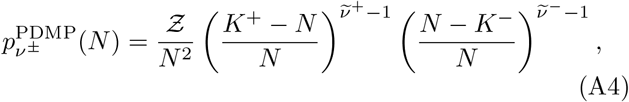

where *Ƶ* is the normalisation constant and *N* ∈ [*K*_−_, *K*_+_]. In addition, to capturing many features of 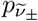, (A4) can be used to accurately approximate the average population size: 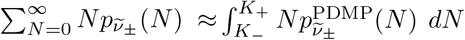 [18, 19, 21]. Eq. (A4) of the binary PSD has also been used to approximately compute the fixation probability and uMFT in two-state switching models [18–21, 23, 27]; see Sec. III B.

It is worth noting that DMN is commonly used to model evolutionary processes because of its simplicity and ability to capture the conditions used in laboratory-controlled experiments. Even if these are generally carried out in periodically changing environments [7, 41, 42], it has been shown that letting *K* switch periodically between *K*^+^ and *K*^−^ with a frequency 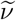 leads to essentially the same dynamics as with Eq. (A1) [21]. Moreover, the relationship between DMN and other forms of environmental noise is well documented [23, 64, 65].

### 2. Multi-state environmental variability

In this work, environmental variability is encoded in the continuous-time Markov process *ξ*(*t*) = *I* ∈ {−*n*, …, 0, …, *n*} that is a (2*n* + 1)-state coloured noise (non-zero correlation time). Here, the environmental noise *ξ*(*t*) is initialised in its stationary distribution (see below) and is therefore a (2*n* + 1)-state generalisation of the classical (two-state) DMN noise used in numerous studies [18–20, 22, 24–28, 41, 43, 63–65].

The generator matrix associated with the multi-state switching of *ξ*(*t*) according to Eq. (4) is the (2*n* + 1) *×* (2*n* + 1) stochastic matrix ***Q*** whose entries *Q*_*j,k*_ are the switching rates of (4) [52]. For convenience, we choose the indices *j, k* = 0, …, 2*n* so that, when *j* ≠ *k, Q*_*j,k*_ corresponds to the rate of the transition from state *j* − *n* to state *k − n*, i.e. 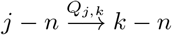. Thus, the generator matrix ***Q*** = (*Q*_*j,k*_) of *ξ*(*t*) defined by the transitions (4) is

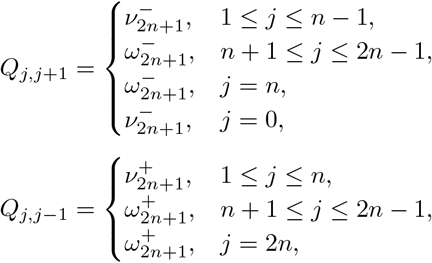

with *Q*_*jj*_ = − ∑_*k≠j*_ *Q*_*jk*_. The stationary distribution of *ξ*(*t*) is the normalised left eigenvector ***π*** of the tridiagonal matrix ***Q*** associated with eigenvalue 0. When 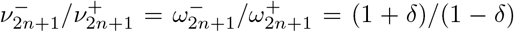, we find that ***π*** = (*π*_*j*_), where *π*_*j*_ ≡ *π*_*i*+*n*_ and

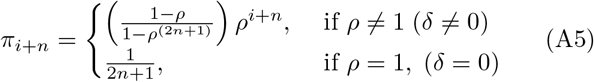

with *ρ* ≡ (1 + *δ*)*/*(1 − *δ*), and *i* = − *n*, …, *n*. For notational simplicity, we denote the stationary distribution of *ξ* by *π*_*i*_; see Eq. (9). The stationary average of *ξ* is

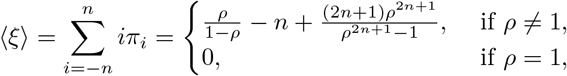

with ⟨*ξ*⟩ *>* 0 if *δ >* 0 and ⟨*ξ*⟩ *<* 0 when *δ <* 0. The stationary distribution ***π*** can also be used to compute the stationary autocovariance *A*(*t*) ≡ lim_*t*_*′*_→∞_ ⟨*ξ*(*t*^*′*^)*ξ*(*t*^*′*^ + *t*) ⟩ − ⟨*ξ*⟩^2^, that decays as 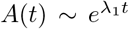, where *λ*_1_ is the eigenvalue of ***Q*** with largest non-zero real part. In general, when *δ* ≠ 0 and *ϵ* ≠ 0, the finite correlation time of *ξ*(*t*) is *t*_*c*_ ∝ 1*/ν*_2*n*+1_ with a proportionality factor that depends non-trivially on *n, δ* and *ϵ*. This is to be compared with the correlation time 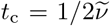 of the dichotomous noise *ξ*_D_ [18, 19, 21, 63–65]; see Appendix A 1.

The multi-state carrying capacity, driven by *ξ*, can be written as

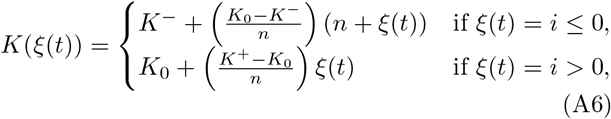

with *K*_*i*_ ≡ *K*(*ξ*(*t*) = *i*) given by Eq. (6), and its stationary probability distribution is therefore also ***π***, with lim_→∞_ ℙ (*K*(*t*) = *K*_*i*_) = *π*_*i*_ given by Eq. (9). The average stationary carrying capacity is therefore 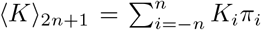 while its stationary variance is 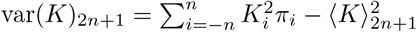.

To meaningfully compare the properties of multi-state and binary switching models, it is useful to compute the mean times for the transitions from *K* = *K*^∓^ to *K* = *K*^*±*^. In binary switching models, where the transitions 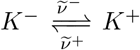 occur at rates 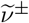, the mean waiting times for the reactions *K*^∓^ → *K*^*±*^ thus are 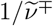. In (2*n* + 1)-switching models, the transition *K*^−^ → *K*^+^ consists of *n* moves, each of mean waiting time 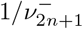, followed by *n* other transitions of average waiting time 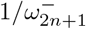; see Fig. 1(a), yielding a mean waiting time 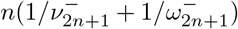. Similarly, the mean waiting time for the transition *K*^+^ → *K*^−^ in multi-state switching models is 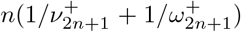. Therefore, by setting 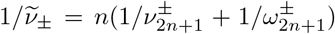, the mean times for the transition between the extreme rates of feast (*K* = *K*^*+*^) and famine (*K* = *K*^−^) is the same in binary and multistate models. With 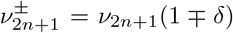 and 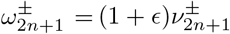, we obtain the expression of Eq. (8).

For the *N*-PDMP-based approximation of Sec. III B 2, we need to compute the average number of switches *N*_sw_ occurring on the evolutionary timescale *t* ~ 1*/s*. This is obtained by multiplying 1*/s* with the total escape rate, given by 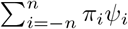, where 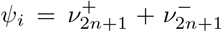 when 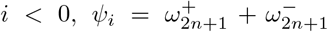 when *i >* 0, and 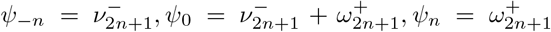 and *π*_*i*_ is given by Eq. (9) (see also Eq. (A5)). The average number of switches during time 1*/s* is therefore 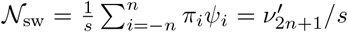. The rescaled switching rate used in the approximations of Eqs. (19) and (B2) is therefore 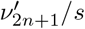, where

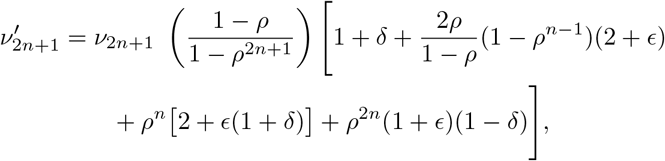

which is the expression of Eq. (18). When *δ* = *ϵ* = 0, we have *ρ* → 1 and this expression simplifies to *ν*^*′*^ = 4*nν*_2*n*+1_*/*(2*n* + 1). Similarly to what was done in Refs. [18–21, 23], we have also tried to use the approximation of Eq. (19) with the simpler rescaled rate *ν*_2*n*+1_*/s* instead of 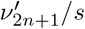, and obtained results that, while in sound agreement with Gillespie simulations, were less accurate than those obtained from Eq. (19) with the effective switching rate given by Eq. (18).

### 3. Master equation

The competition dynamics in the (2*n*+1)-state switching models is a continuous-time multivariate Markov process – defined by the transition rates 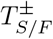 given by Eq. (2) that satisfies the master equation for the joint probability 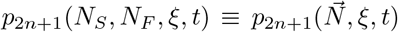 of finding the population in configuration 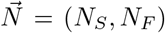 and environmental state *ξ* = *i* ∈ {−*n*, …, *n*}. The master equation for (2*n* + 1)-state switching model reads

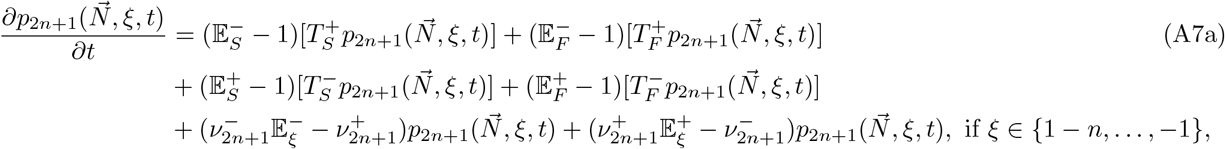

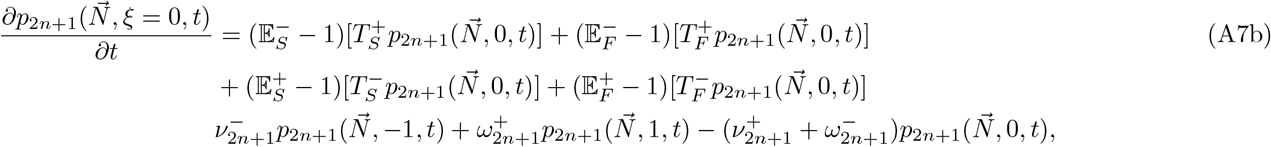

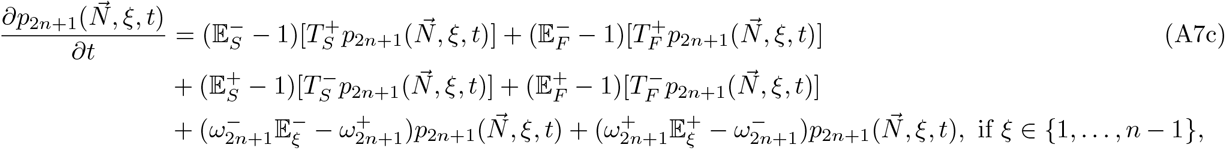

where 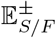 and 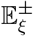 are shift operators such that 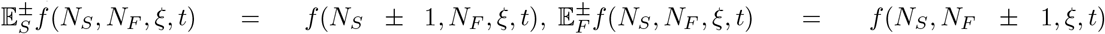 and 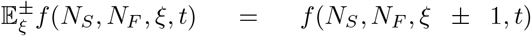, with 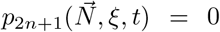 whenever *N*_*S*_ *<* 0 or *N*_*F*_ *<* 0, or *ξ* ∈*/* {−*n*, …, *n*}. The last lines of (A7a)-(A7c) encode environmental switching respectively across harsh/mild states, and through the median state (see Sec. II A). The transition rates are as in Eq. (2), and every time there is a switch *ξ* = *i* ↔ *ξ* = *i* + 1, the carrying capacity appearing in 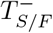 switches *K*_*i*_ ↔ *K*_*i*+1_ according to Eq. (5).

The class of individual-based multi-state switching models defined by Eqs. (2)-(6) satisfies the master equation (A7), and has been simulated using the Gillespie algorithm [66]; see Appendix C.

In Fig. 2, we show the histograms of the marginal quasi-stationary PSD 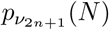 for *t >* 1*/s* obtained by marginalising the joint probability with respect to the environmental states and population composition, i.e. 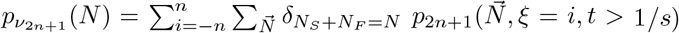, where 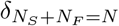 is the Kronecker delta function ensuring that the population size is *N*.

## Appendix B Unconditional mean fixation time of in static and fluctuating environments

In this appendix, we discuss the properties of the mean time for either *S* or *F* to take over the entire population, i.e. the unconditional mean fixation time (uMFT). We first compute the uMFT in a static environment, with a constant carrying capacity 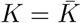, using an approximation based on the Moran process. We then use this result to study the uMFT in fluctuating environments with a multi-state switching carrying capacity.

### 1. uMFT in static environment: the Moran approximation

The uMFT can be computed exactly for a population of constant size 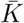 evolving according to the classical Moran process [8, 53]; see Sec. II C. When the initial fraction of *S* individuals is 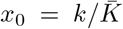, its expression, here denoted by *τ*_*M*_, defined as the expected value of the unconditional fixation times 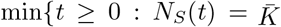 or *N*_*S*_(*t*) = 0}, reads [21, 25, 27, 55, 56, 71]

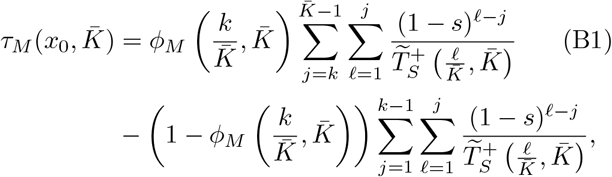

where 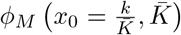 is given by Eq. (12) and 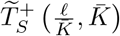 is from Eq. (11) with 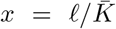. When 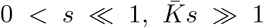, with *x*_0_ = *O*(1) (weak selection, large population, and initial condition well separated from the absorbing boundaries 0 and 1), we have 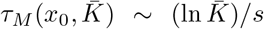: The uMFT has a weak dependence on 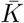, with fixation occurring on a timescale 1*/s* [8, 18–20, 27, 54].

The Moran uMFT, given by Eq. (B1), is a good approximation of the uMFT in a static environment with a constant carrying capacity, 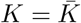, where the total population size fluctuates about 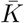, i.e. 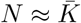. In this case, when 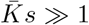 and 0 *< s* ≪ 1 (and *x*_0_ = *O* (1)), fixation occurs on a timescale 1*/s* [12, 13, 18, 19, 21, 24–28].

### 2. uMFT under environmental switching

Similarly to what is done in Sec. III B for the *S* fixation probability, the Moran uMFT, given by Eq. (B1), can be used to obtain suitable approximations of the uMFT under (2*n* + 1)-state environmental switching, denoted by *τ*_2*n*+1_ and defined as the expected value of min {*t* ≥ 0 : *N*_*S*_(*t*) = *N*(*t*) *>* 0 or *N*_*F*_ (*t*) = *N*(*t*) *>* 0. To study *τ*_2*n*+1_, it is again convenient to distinguish the regimes *ν*_2*n*+1_ ≪ *s, ν*_2*n*+1_ ≫ *s*, and *ν*_2*n*+1_ ~ *s*. Since the uMFT in static environments scales with 1*/s* for weak selection and a sufficiently large population, we expect a similar behaviour in fluctuating environments for all *n*, when 0 *< s* ≪ 1, *s* ⟨*N*⟩ *>* 1, and *x*_0_ = (1), as in the binary case [18, 19, 21, 27].

In the slow switching regime (*ν*_2*n*+1_ ≪ *s*), it is likely that no environmental switches occur prior to fixation. In this regime, a Moran approximation for the uMFT is obtained from Eq. (B1) for a population of size *N* ≈ *K*_*i*_ with a probability *π*_*i*_, yielding 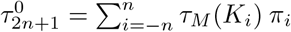 (compare with Eq. (16)). For notational simplicity, we have dropped the dependence on *x*_0_.

Under fast environmental switching (*ν*_2*n*+1_ ≫ *s*), the carrying capacity self-averages and the population experiences the effective carrying capacity *K*_2*n*+1_ given by Eq. (13). A Moran approximation of the uMFT is thus given by Eq. (B1) for a population of size *N* ≈ *K*_2*n*+1_, yielding 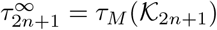 (compare with Eq. (17)).

In the intermediate switching regime (*ν*_2*n*+1_ ~ *s*), when *s* ≪ 1, we can use the *N*-PDMP-based approximation employed in Sec. III B; see Eq. (19). Exploiting the timescale separation between *N* and *x*, and using the *N*-PDMP density (see Eq. (14)) to approximate the stationary PSD, we can write 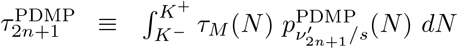, using Eq. (18) [18–21, 27].

The uMFT in the fluctuating multi-state switching models with (2*n* + 1) environmental states can therefore be approximated by

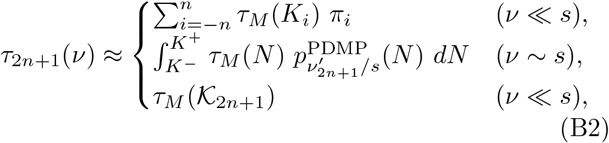

using Eqs. (B1), (9), (6), and (13). Simulation results of Figs. 7 and 8 show that Eq. (B2) is generally a good approximation as it faithfully captures the main features of the uMFT across all switching regimes. As expected, the uMFT is found to scale as 1*/s*, with 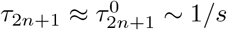 when *ν*_2*n*+1_ ≪ *s* and 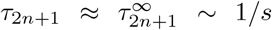 when *ν*_2*n*+1_ ≫ *s*, as shown by the dotted and dashed lines. Moreover, 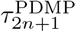 correctly predicts that the uMFT is a decreasing function of *ν*_2*n*+1_ (all other parameters kept fixed) in Figs. 7 and 8, and we find 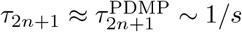 in all switching regimes. The uMFT at a given *ν*_2*n*+1_ increases with *n* (all other parameters being kept fixed); compare, e.g., Figs. 7(a) and 8(a). Similarly, at a given ν_2n+1_, τ_2n+1_ increases with δ; see, e.g., Fig. 7(b). Simulation results in Figs. 7, 8 and Fig.10(b) show no noticeable dependence on the parameter ϵ, when −0.8 ≤ *ϵ* ≤ 5.

**FIG. 7.**
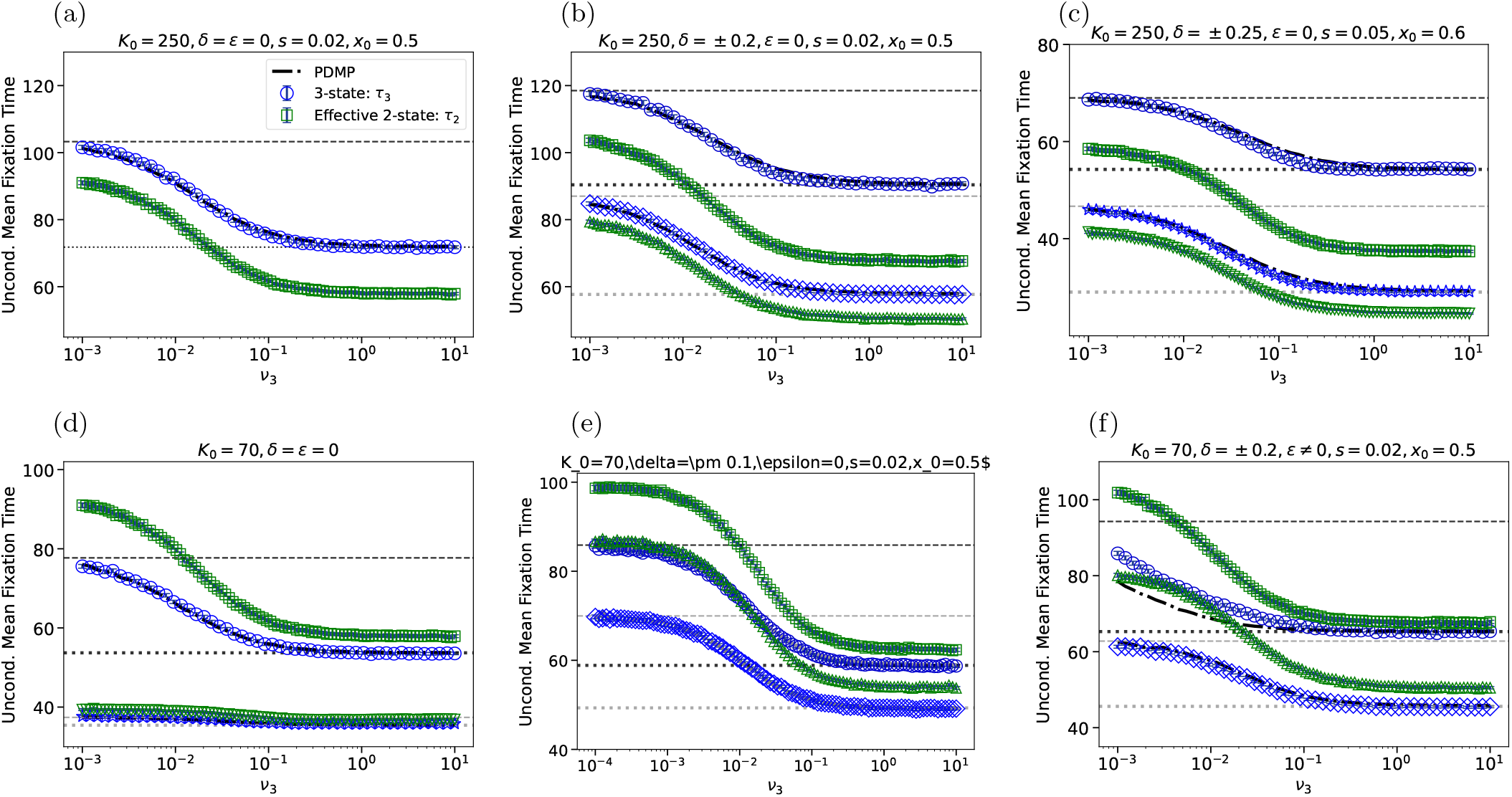
Unconditional mean fixation time in the three-state switching model as a function of *ν*_3_. The centre state carrying capacity is *K*_0_ = 250 in (a)-(c), while *K*_0_ = 70 in (d)-(f). Here, *n* = 1, and (*K*^−^, *K*^+^) = (50, 450) in all panels except (c) where (*K*^−^, *K*^+^) = (25, 475). Symbols are from full stochastic simulations and black dashed-dotted curves (in all panels), often almost indistinguishable from markers, are from the *N*-PDMP-based approximation; see Eq. (19). In blue, is shown *τ*_3_ vs. *ν*_3_ for different parameter sets (*δ, ϵ, s, x*_0_) that are the same as in Fig. 5 (see also the legends). Green symbols show simulation data for *τ*_2_, the uMFT of the effective two-state model; see text and Eq. (8). In (b,c,e,f), results for *τ*_2_ are shown as triangles when *δ* = v 0.2 and squares when *δ* = −0.2. In (d), simulations data of *τ*_2_ are shown as squares for (*s, x*_0_) = (0.02, 0.5) and downside triangles for (*s, x*_0_) = (0.1, 0.7). For the *N*-PDMP-based approximation of *τ*_2_, see Refs. [18, 19, 21]. The horizontal dashed and dotted lines are eyeguides showing 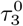 (dashed) and 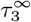 (dotted). Error bars are included but typically too small to see.

**FIG. 8.**
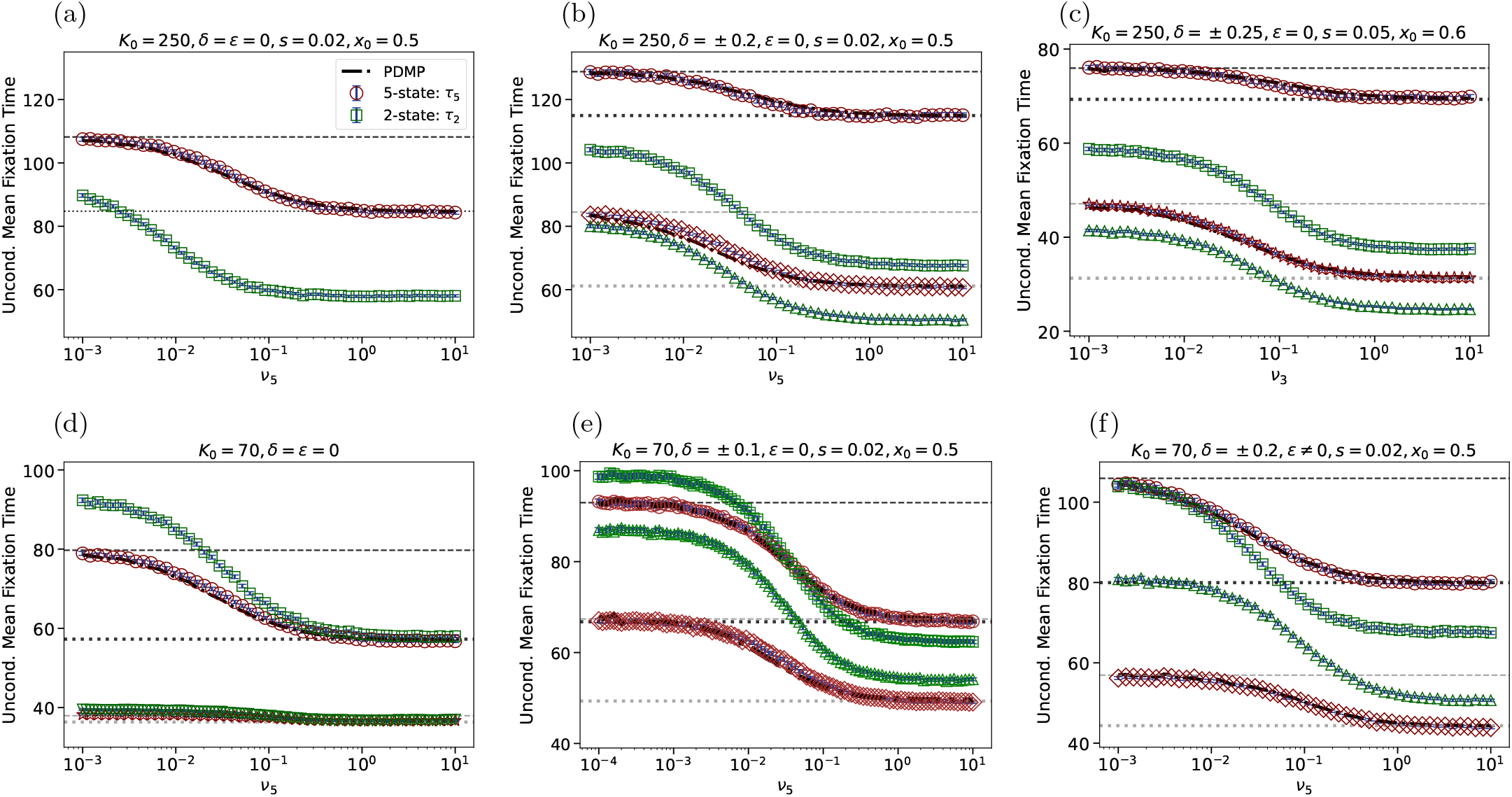
Unconditional mean fixation time in the five-state switching model as a function of *ν*_5_. The centre state carrying capacity is *K*_0_ = 250 in (a)-(c), while *K*_0_ = 70 in (d)-(f). Here, *n* = 1, and (*K*^−^, *K*^+^) = (50, 450) in all panels except (c) where (*K*^−^, *K*^+^) = (25, 475). Symbols are from full stochastic simulations and dashed-dotted curves (in all panels), almost indistinguishable from markers, are from the *N*-PDMP-based approximation; see Eq. (19). In brown, is shown *τ*_5_ vs. *ν*_5_ for different parameter sets (*δ, ϵ, s, x*_0_) that are the same as in Fig. 6 (see also the legends). Green symbols show simulation data for *τ*_2_, the uMFT of the effective two-state model; see text and Eq. (8). In (b,c,e,f), results for *τ*_2_ are shown as triangles when *δ* = −0.2 and squares when *δ* = − 0.2. In (d), simulations data of *τ*_2_ are shown as squares for (*s, x*_0_) = (0.02, 0.5) and downside triangles for (*s, x*_0_) = (0.1, 0.7). For the *N*-PDMP-based approximation of *τ*_2_, see Refs. [18, 19, 21]. The horizontal dashed and dotted lines are eyeguides showing 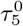 (dashed) and 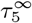 (dotted); see Eq.(B2). Error bars are too small to see.

**FIG. 9.**
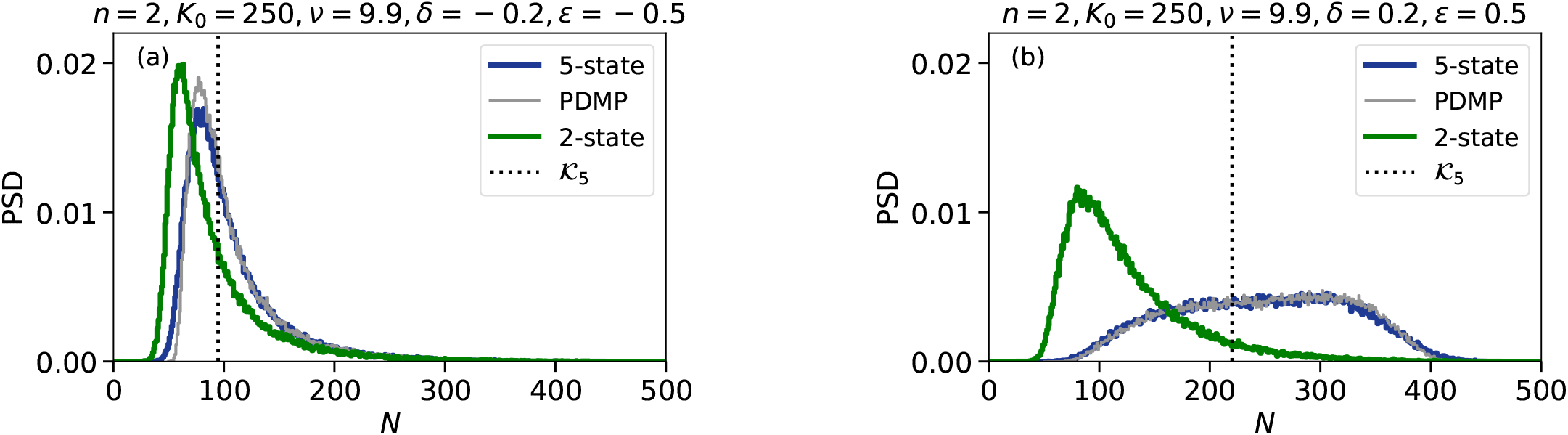
Supplementary figure showing in dark blue the PSD of the five-state switching model 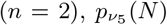 for *ν*_5_ = 9.9 (fast switching), with (*s, x*_0_, *K*^−^, *K*_0_, *K*^+^) = (0.02, 0.5, 50, 250, 450). Other parameters are: (*δ, ϵ*) = (0.2, 0.5) in (a) and (*δ, ϵ*) = (− 0.2, − 0.5) in (b). In grey is shown the *N*-PDMP approximation of the PSD (see Eq. (14)), while the green histograms are from the effective two-state model (see text). Dotted lines are eyeguides indicating *K*_5_ ≈ 94.70 (a) and *K*_5_ ≈ 220.18 (b). These histograms have been obtained for *t >* 2000 (Appendix C) and have to be compared with those of Fig. 2(g),(h).

**FIG. 10.**
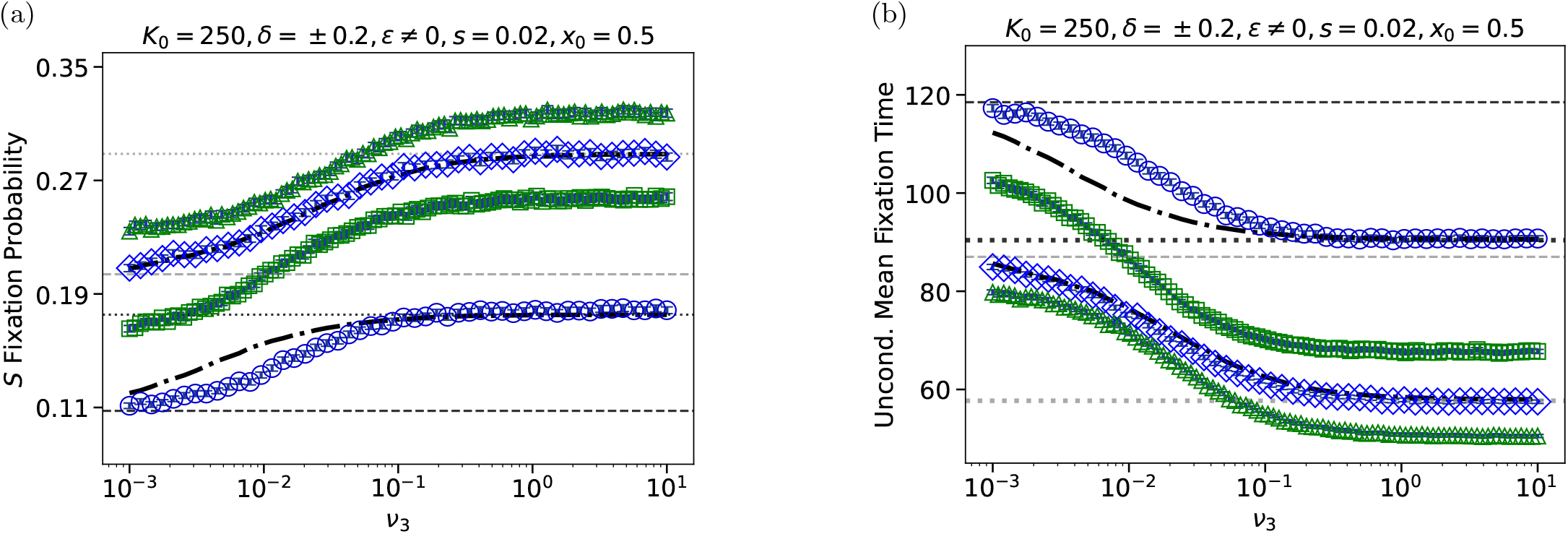
Supplementary figure showing in blue *ϕ*_3_ and *τ*_3_ vs *ν*_3_ under ternary switching for (*δ, ϵ*) = (0.2, 5) (circles) and (*δ, ϵ*) = (−0.2, −0.5) (diamonds). Other parameters are (*s, x*_0_, *n, K*^−^, *K*_0_, *K*^+^) = (0.02, 0.5, 1, 50, 250, 450). The dashed-dotted black curves show the *N*-PDMP approximations 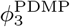 in (a) and 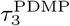 in (b); see Eqs. (19) and (B2). For (*δ, ϵ*) = (−0.2, −0.5), 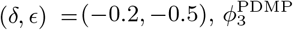 and 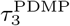 are almost indistinguishable from *ϕ*_3_ and *τ*_3_. The green symbols show *ϕ*_2_ in (a) and *τ*_2_ in (b) for (*δ, ϵ*) = (0.2, 5) (squares) and (*δ, ϵ*) = (−0.2, −0.5) (triangles).

## Appendix C Simulation methods

To complement our analytical approaches, we have performed extensive stochastic simulations of the multistate switching models using the Gillespie algorithm [66]. These exactly mirrors the continuous-time dynamics encoded in the master equation (A7). The implementation of the simulations was efficiently optimised through just-in-time compilation (Numba) [81]. In Sections III B 3 and III B 4, we have specifically focused on the switching models with three (*n* = 1) and five (*n* = 2) environmental states, whose PSD, average population size, and fixation properties were computed; see Figs. 2 and 3, and Figs. 5-10. The PSD, *S* fixation probability and uMFT have been compared with their binary-state counterparts [18, 19, 21]. The multi-state environmental noise *ξ* and carrying capacity *K*; see Eqs. (3)-(6) and (A6), are always at stationarity, and therefore each simulation starts a time *t* = *t*_0_ = 0 with an initial value of *ξ*(*t*_0_) and carrying capacity *K*(*t*_0_) drawn from their stationary distribution ***π***; see Eq. (9). In each run, the population has an initial size *N*(*t*_0_) = ⟨*K*⟩_2*n*+1_ that coincides with the average carrying capacity, and its composition is *N*_*S*_(*t*_0_) = round (*x*_0_*N*(*t*_0_)) and *N*_*F*_ (*t*_0_) = *N*(*t*_0_) − *N*_*S*_(*t*_0_), where *x*_0_*N*(*t*_0_) is rounded to the nearest integer. and Simulations were run in batches of 10^5^ realizations for each set of parameters {*n, K*^−^, *K*^+^, *K*_0_, *s, x*_0_, *ν*_2*n*+1_, *δ, ϵ*} by accounting for all the possible reactions that can take place at each time increment. For the class of individualbased models studied here, these are the four possible birth or death reactions with rates 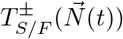; see Eq. (2), and the switching of the multi-state carrying capacity *K*(*t*) driven by the environmental switches of *ξ*.

In Figs. 5-8 and 10, for each set {*n, K*^−^, *K*^+^, *K*_0_, *δ, ϵ*}, simulation data for *ϕ*_3_, *ϕ*_5_, *τ*_3_ and *τ*_5_ are shown for 50 values of *ν*_3,5_ ∈ [10^−3^, 10] in all panels, excepts in panels (e) of these figures consisting of 100 values of *ν*_3,5_ ∈ [10^−4^, 10]. The panels (a)-(d),(f) of Figs. 5 and 7 also contain simulation results for *ϕ*_2_ and *τ*_2_ obtained for 100 values of *ν*_3_ ∈ [10^−3^, 10] in panels (a)-(d),(f), and for 100 values of *ν*_3_ ∈ [10^−4^, 10] in Fig. 5(e) and 7(e). The panels (a)-(d),(f) of Figs. 6 and 8 contain simulation data of *ϕ*_2_ and *τ*_2_ obtained for 50 values of *ν*_5_ ∈ [10^−3^, 10] in panels (a)-(d),(f), and for 100 values of *ν*_5_ ∈ [10^−4^, 10] in Fig. 6(e) and 8(e). Each data point in Figs. 2 and 3, and Figs. 5-10 has been obtained by sampling ℛ = 10^5^ realizations. The statistical errors on the simulation results on the fixation probability *ϕ*_2*n*+1_ and *ϕ*_2_ of Figs. 5, 6 and 10 has been estimated using the classical Wald confidence interval method. Accordingly, the estimated error on sample-averaged value 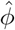 of *ϕ* is 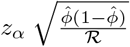, where we have chosen *z*_*α*_ = 1.96, which corresponds to a 95% confidence interval. For ℛ = 10^5^, this yields estimated errors of order 2 · 10^−3^, i.e around 1% to 2% of 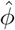 in Figs. 5, 6 and 10. These statistical errors are almost unnoticeable, suggesting that the statistical results are robust.

The histograms *p*_*m*_(*N*), with *m* 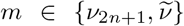 and 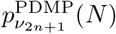 of Figs. 2 and 9 have been obtained by sampling a long trajectory of *N*(*t*) and *N*^PDMP^(*t*) after discarding an initial transient, i.e. for *t > t*_burn_ = 2000 to ensure that *N*(*t*) and *N*^PDMP^(*t*) have settled in their long-time dynamics. For the full individual-based switching models, the long trajectory *N*(*t*) is obtained from Gillespie simulations. For the *N*-PDMP, a long trajectory *N*^PDMP^(*t*) is generated by implementing the environmental switching according to the Gillespie algorithm and, between each switch, by solving the logistic equation 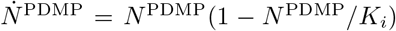 in the current environmental state *i*; see Eq. (14). The histograms representing *p*_*m*_ were thus obtained by taking *ℛ* ≫ 1 samples each spaced by 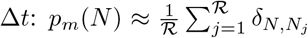, where *N*_*j*_ = *N*(*t*_burn_ + *j*Δ*t*), and 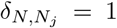 when *N* = *N*_*j*_ and 0 otherwise. Similarly, the stationary *N*-PDMP density 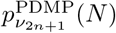 is approximated numerically from a long *N*-PDMP trajectory (after discarding an initial transient *t*_burn_) by collecting a large number ℛ of samples, each spaced by Δ*t*. These define the empirical measure 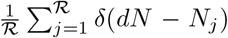, where *δ*() is a Dirac delta function. Computationally, the sampled values are grouped into bins of unit width, and the resulting counts are normalised to construct the histogram approximations of the stationary *N*-PDMP density shown in Fig. 2. For each sampled trajectory, the total simulated time is therefore *t*_burn_ + ℛΔ*t*. In Figs. 2 and 3, we have used ℛ = 10^5^ and Δ*t* = 10. Within this approach, the average population size of Fig. 3 was computed as the discrete time average from a long trajectory of *N*(*t > t*_burn_) and *N*^PDMP^(*t > t*_burn_) according to 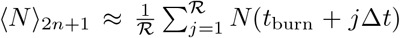 and 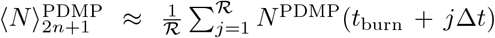, with *t*_burn_ = 5000. The estimated error on ⟨*N*⟩ decreases with the number *ℛ*_ind_ of independent samples, as 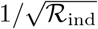. This implies that this approach is well-suited to efficiently compute the average population size when the switching rate is not too low. In Fig. 3, for *ℛ* = 10^5^ and Δ*t* = 10, the estimated error scales as 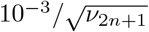 and hence decreases by a factor 100 when *ν*_2*n*+1_ varies from 0.01 to 100. In the limit *ν*_2*n*+1_ → 0, the average population size is given by ⟨*N*⟩_2*n*+1_ ≈ ⟨*K*⟩_2*n*+1_; see Fig. 3 (dotted lines)

The number of independent samples ℛ_ind_ is large when ℛ ≫ 1 and the switching rate is not too low. Therefore, within the approach outlined above, the *N*-PDMP-based approximation of the fixation probability and uMFT are efficiently computed by averaging over ℛsamples of a *N*−PDMP trajectory *N*^PDMP^(*t > t*_burn_) obtained for the rescaled switching rate *ν*^*′*^*/s* (see Eq. (18)) according to 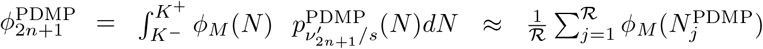 and 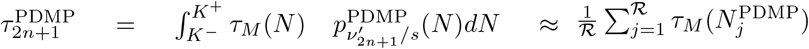, where 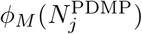 and 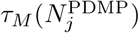 are the Moran fixation probability and uMFT (given by Eqs. (12) and (B1)) evaluated at 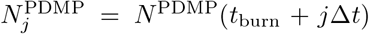. In Figs. 5-8 and Fig. 10, this method was used for *ν*^*′*^*/s* ~ 0.1 − 1000, with ℛ = 10^5^, Δ*t* = 10 and *t*_burn_ = 2000, and proved to be fast and reliable: It provided results that are generally in very good agreement with those of Gillespie simulations (and analytical computations), but obtained much faster (by a factor of up to 100).

## Appendix D Supplementary figures

To analyse a possible dependence of the PSD, *S* fixation probability and uMFT on the parameter *ϵ*, we have reproduced the histograms of Fig. 2 and computed *ϕ*_2*n*+1_ and *τ*_2*n*+1_ for different values of *ϵ*, with − 0.8 ≤ *ϵ* ≤ 5, and obtained results that are essentially the same as those in Fig. 2 and Figs. 5-8 where *ϵ* = 0. This is illustrated in Fig. 9(a),(b) where we have used the same parameters as in Fig. 2(g),(h), but now set *ϵ* = *±*0.5 rather than *ϵ* = 0. In Fig. 9(c),(d) we also show the simulation data of *ϕ*_3_ and *τ*_3_ for the same parameters as in Figs. 5(b) and 7(b), but now with *ϵ* = −0.5 and *ϵ* = 5 instead of *ϵ* = 0. The histograms of Fig. 9(a),(b) are identical to those of Fig. 2(g),(h), and the results for *ϕ*_3_ and *τ*_3_ in Fig. 9(a),(b) are essentially the same as in Fig. 5(b) and Fig. 7(b). This suggests that the PSD, *ϕ*_2*n*+1_ and *τ*_2*n*+1_ have no noticeable dependence on the parameter *ϵ* for −0.8 ≤ *ϵ* ≤ 5.

